# No evidence for whole-chromosome dosage compensation or global transcriptomic expression differences in spontaneously-aneuploid mutation accumulation lines of *Saccharomyces cerevisiae*

**DOI:** 10.1101/2020.12.01.404830

**Authors:** Holly C. McQueary, Megan G. Behringer, Sam Demario, Alexander Joao Jamarillo Canas, Brittania Johnson, Ariella Tsfoni, John Chamberlin, David W. Hall

**Affiliations:** University of Georgia; Vanderbilt University; University of California Los Angeles

**Keywords:** Aneuploidy, dosage compensation, gene expression, Mutation accumulation

## Abstract

Aneuploidy, the state in which an organism’s genome contains one or more missing or additional chromosomes, often causes widespread genotypic and phenotypic effects. Most often, aneuploidies are deleterious; the most common examples in humans being Down’s syndrome (Trisomy 21) and Turner’s syndrome (monosomy X). However, aneuploidy is surprisingly common in wild yeast populations. In recent years, there has been debate as to whether yeast contain an innate dosage compensation response that operates at the gene, chromosome, or the whole-genome level, or if natural isolates are robust to aneuploidy without such a mechanism. In this study, we tested for differential gene expression in 20 aneuploid and 16 euploid lines of yeast from two previous mutation accumulation experiments, where selection was minimized and therefore aneuploidies arose spontaneously. We found no evidence for whole-chromosome dosage compensation in aneuploid yeast but did find some evidence for attenuation of expression on a gene-by-gene basis. We additionally found that aneuploidy has no effect on the expression of the rest of the genome (i.e. “trans” genes), and that very few mutually exclusive aneuploid lines shared differentially expressed genes. However, we found there was a small set of genes that exhibited a shared expression response in the euploid lines, suggesting an effect of mutation accumulation on gene expression. Our findings contribute to our understanding of aneuploidy in yeast and support the hypothesis that there is no innate dosage compensation mechanism at the whole-chromosome level.

## Introduction

Aneuploidy occurs when an organism contains an abnormal number of one or a few chromosomes. Familiar examples are those causing human disorders, such as Down’s syndrome (trisomy 21) or Turner’s syndrome (monosomy X) (Hassold and Hunt 2001). While autosomal aneuploidies are generally deleterious in most organisms, presumably because of dosage problems (Chunduri and Storchova 2019), in some species, aneuploidies are surprisingly common, such as in some wild yeast (*Saccharomyces cerevisiae*) isolates (Strope *et al.* 2015). It has been shown experimentally that the accumulation or loss of chromosomes can be adaptive in certain environments (Selmecki *et al.* 2006; Pavelka *et al.* 2010; Chen *et al.* 2012; Yona *et al.* 2012; Selmecki *et al.* 2015; de Vries *et al.* 2018). For example, yeast grown in an oxide-rich medium accumulate an extra copy of chromosome XI as a response to oxygen stress (Kaya *et al.* 2015), and resistance to fluconazole in *Candida albicans* often involves an increase in copy number of a single chromosome (Wakabayashi *et al.* 2017; Koo *et al.* 2018). There is debate as to how aneuploidy is tolerated in wild populations. Some hypothesize there is an intrinsic mechanism of transcriptional dosage compensation to buffer the deleterious effects of imbalanced gene dosage (Hose *et al.* 2015; Gasch *et al.* 2016), similar to the mechanism of dosage compensation observed in sex chromosomes (Marin *et al.* 2000). Such autosomal compensation has been observed in *Drosophila* and other species (Birchler *et al.* 1990; Matos *et al.* 2015). In yeast, the presence of such a mechanism has been debated, with some studies concluding that there is no evidence for transcriptional dosage compensation at the whole-chromosome level (Torres *et al.* 2010), and others suggesting that aneuploid wild yeast post-translationally attenuate protein levels by increasing protease activity or upregulating genes that are part of multiprotein complexes so that the relative dosages are more even (Chen *et al.* 2003; Veitia *et al.* 2008).

While dosage compensation has been observed for autosomes in *Drosophila* (Devlin *et al.* 1982; Birchler *et al.* 1990; McAnally and Yampolsky 2009; Chen and Oliver 2015; Hangnoh Lee 2016; Lee *et al.* 2016), it is unknown whether such an intrinsic mechanism exists in yeast. The fact that yeast are often found to be aneuploid in natural isolates (Strope *et al.* 2015) and develop aneuploidies in response to certain environmental conditions (Selmecki *et al.* 2006; Pavelka *et al.* 2010; Chen *et al.* 2012; Yona *et al.* 2012; Selmecki *et al.* 2015; de Vries *et al.* 2018) could suggest that aneuploidy causes changes in gene expression that are adaptive, and no DC exists (Kaya *et al.* 2015; Linder *et al.* 2017). Alternatively, yeast may be naturally robust to aneuploidy, so that aneuploid strains do not differ in fitness and thus occur in nature as neutral variants. The second hypothesis, coupled with the occurrence of aneuploid strains at reasonably high frequencies, suggests that yeast may contain an innate mechanism for attenuating or compensating for differences in gene dose and that mutation to aneuploidy is relatively frequent.

Understanding dosage compensation is important for several reasons. Aneuploidy cannot be avoided because segregation machinery is not perfect, and mitotic and meiotic cells both experience nondisjunction events. As such, determining whether there are intrinsic mechanisms of dosage compensation gives insight into the likely consequences of such aneuploidy. Dosage compensation is also critically important during the evolution of sex chromosomes from homomorphic autosomes (Charlesworth 1991). Dosage compensation is thought to play a critical role in the evolution of sex chromosomes because of their different copy numbers in males versus females, a common example being the X-chromosome in XY systems. There are a variety of ways in which dosage compensation occurs in sex chromosomes to make up for differences in gene dosage between the sexes (Chandler 2017). However, the degree to which compensation evolves prior to, during, or after the evolution of dimorphism remains an open question (Gu and Walters 2017). More information on the consequences of aneuploidy in spontaneously-aneuploid yeast genomes can provide further insights into the evolution of sex chromosomes and imbalances in gene dose.

To fully understand the effects of aneuploidy on yeast populations, we seek estimates of the rate of aneuploidy and the effects of aneuploidy on gene expression. Previous studies have observed the effects of aneuploidy in wild yeast populations, where selection is acting; and in chemically-or mitotically-induced aneuploids, where the rate of production of aneuploids is being manipulated (Linder *et al.* 2017); (Campbell *et al.* 1981; Anders *et al.* 2009; Mulla *et al.* 2014). In this study, we sought to determine the spontaneous rate of aneuploid formation for each chromosome and the effects of aneuploidy on gene expression in two strains of diploid yeast *in the absence of selection*. In each strain, spontaneous aneuploid events were captured during a 2000-generation mutation accumulation (MA) experiment with a single-cell bottleneck every 20 generations (joseph and hall 2004a; Zhu *et al.* 2014). By passaging through a single-cell bottleneck the effective population size is kept small (*N*_e_ ≈ 11), which minimizes the effects of selection; only mutations with heterozygous fitness effects (*s*) of approximately 5% or greater (i.e. 2*s* ≥ 1/11) will be efficiently acted on by selection (Wright 1931). Using RNA sequencing, we analyzed the gene expression of 20 aneuploid and 16 euploid lines across both strains to find differentially expressed genes and to determine if there was evidence for dosage compensation at the whole-chromosome and individual-gene levels in yeast.

## Materials and Methods

### Estimating the spontaneous rate of aneuploid mutation

To determine the rate at which spontaneous aneuploidy occurs in yeast, we analyzed data from two previous mutation accumulation (MA) experiments (Joseph and Hall 2004b). In both, an ancestral strain was copied into multiple MA lines, which were then maintained separately for ~2000 cell generations (*G*) (2063 generations in the homozygous ancestor lines and 2108 in the heterozygous ancestor lines) via single-colony transfer every 48 hours (± 1 hour) for 100 transfers on solid YPD (1% yeast extract, 2% peptone, 2% glucose, 2% agarose) medium. The actual number of generations that passed was estimated by measuring colony size after 48 hours of growth in a representative sample of lines and passages and then determining cell number by counting using a hemocytometer. To confirm that the vast majority of cells present at 48 hours were viable, we also estimated cell number by serial dilution and plating (data not shown).

The two diploid ancestral strains differed in their origin and degree of heterozygosity. One strain was obtained from a mating between NCYC 3631, which is a Matαderivative of YPS606 (an oak strain from Pennsylvania, USA), and NCYC3596, a Mat*a* derivative of DBPVG1106 (a wine strain isolated from a lici fruit in Indonesia). This highly heterozygous strain had a heterozygous site every ~250 bp and was homozygous for *ho* and *ura3* mutations.

The other strain was derived from a standard lab strain (S228C) and carried the following mutations: *ho ade2, lys2-801, his3-ΔD200, leu2-3.112,* and *ura 3-52* (Joseph and Hall 2004a). The strain was obtained by transforming a Mat*a* haploid version of the strain with an *HO URA3* plasmid to generate a diploid version of the strain, followed by counterselection of the plasmid on 5FOA (Joseph and Hall 2004a). This strain was thus homozygous at all loci except the mating type locus.

We used the number of aneuploid chromosomes in the MA lines at the end of the experiment to calculate the rate at which aneuploidy occurs in each of these strains. In brief, if the rate of aneuploidy for chromosome *c* is *μ*_*c*_, then the probability that a line is not aneuploid for this chromosome is (1-*μ*_*c*_)^G^, where G is the number of generations of MA. Thus if *nc* MA lines show aneuploidy for this chromosome, implying that (*n - n*_*c*_) do not, where *n* is the total number of MA lines, then we can estimate the rate of aneuploidy per chromosome by solving (1-*μ*_*c*_)^G^ = (1 - *n*_*c*_ /*n*) for *μ*_*c*_. Similarly, we can estimate the overall aneuploidy rate, *μ*, which is the probability that a cell will become aneuploid for any chromosome in a single cell division, by solving (1-*μ*)^G^ = (1 – *n*_*a*_ /(16 *n*)) for *μ*, where *n*_*a*_ is the number of aneuploid chromosomes across all MA lines.

### Estimating the effects of aneuploidy on gene expression

To determine the effects of aneuploidy on gene expression, we collected and analyzed RNA sequencing data from a selection of euploid and aneuploid lines from each experiment. For aneuploid samples, we chose all the MA lines that were monosomic for a chromosome (3 lines), those that shared common aneuploidies (21 lines), and those that had more than one aneuploidy event (4 lines). From the homozygous ancestor experiment, we selected 10 aneuploid and 12 euploid MA lines. From the heterozygous ancestor experiment, we selected 10 aneuploid and 6 euploid MA lines. Additionally, we collected RNA sequencing data for both ancestral lines. The MA and the ancestral lines that had been stored at −80°C were pulled from the freezer by streaking onto YPD plates (1% yeast extract, 2% peptone, 2% glucose, 2% agarose) for RNA extraction (see below). The homozygous strains were run in two separate RNA sequencing runs, separated by 2 years. In both sequencing runs, we included three replicates of each ancestor. For analysis, we kept these two datasets separate because we found that the ancestor was significantly different across the two sequencing runs. Across both strains and sequencing runs, we obtained RNA sequencing data for 38 strains representing two ancestor strains, 20 euploid lines and 16 aneuploid lines.

For obtaining RNA, each line was pulled onto solid YPD and allowed to grow for two days at 30°C, and then three individual colonies (biological replicates) were used to inoculate three 3ml liquid YPD cultures (no agarose) of each line. Liquid cultures were incubated on a rotator at 30°C for 24 hours, before being diluted into 50ml YPD and allowed to grow on a shaker at 30°C for 6 hours. Optical density (OD) measurements were taken to ensure all cultures were in the same log growth phase. Cells were then pelleted, and RNA was extracted from each replicate using the MasterPure Yeast RNA Purification Kit (Epicentre). Integrity, concentration, and quality of RNA samples were assessed using a Qubit (Thermo Fisher Scientific). Libraries were prepared using the Illumina Stranded RNAseq Kit and were sequenced at the Georgia Genomics and Bioinformatics Core (https://dna.uga.edu/) on the Illumina NextSeq (75 cycles) single end 75bp reads High Output flow cell. Samples were multiplexed and split across two sequencing lanes.

Raw reads were processed by the Georgia Genomics and Bioinformatics Core to remove sequencing adapters and demultiplex samples. Quality control was performed using FastQC version 1.8.0_20 with default parameters (available at www.bioinformatics.babraham.ac.uk/projects/fastqc/). Low-quality bases were trimmed using Trimgalore version 0.4.4 using -phred 33, -q 20 (available at www.bioinformatics.babraham.ac.uk/projects/trim_galore/). RNA sequences were aligned to the *Saccharomyces cerevisiae* reference genome (UCSC version sacCer3, available at support.illumina.com/sequencing/sequencing_software/igenome.html) and transcripts were annotated using Tophat v. 2.1.1 with -i 10 -I 10000 (Trapnell *et al.* 2012). Cufflinks v. 2.2.1 was used to assemble sample transcriptomes using default parameters (Trapnell *et al.* 2012) and Cuffnorm v. 2.2.1 was used with default parameters to normalize reads. Differential expression was determined using Cuffdiff v. 2.2.1 with default parameters, and raw read counts were found using HTseq v. 0.6.1pl (Python v. 2.7.8) (Anders *et al.* 2015). Finally, we used Samtools v. 1.3.1 to convert *.sam* files into *.bam* files and sort the resulting *.bam* files (Li *et al.* 2009). Bash scripts for genome assembly and annotation can be found at https://github.com/hollygene/Dosage_Compensation/tree/master/src/bash_scripts.

To compare chromosome-level changes in gene expression across strains, Cuffnorm v. 2.2.1 (Trapnell *et al.* 2012) was used to calculate FPKM (fragments per kilobase per million reads) for each RNAseq data set. A custom bash script was then generated to join the FPKM values for each strain with the gene annotations file, convert the resulting file into a *.csv* formatted file, remove mitochondrial sequences (as we were not interested in mitochondrial gene expression), and change the chromosome names from Roman numerals to numbers (script can be found at https://github.com/hollygene/Dosage_Compensation/blob/master/src/bash_scripts/old/DC_workflow_April2017.sh). For each gene, the average FPKM across the three replicates for each strain was calculated, followed by the average FPKM ratio (average FPKM in an MA line divided by the average FPKM in the ancestor). We noticed that the FPKM ratio was highly variable across MA line replicates for genes with an average FPKM < 5 across all euploid strains (ancestor + euploid MA lines), so we removed such genes, leaving 6181 genes. We also removed rRNA genes and tRNA genes, as these are challenging to map accurately and, because of their propensity to show extreme variation in copy number, can cause issues with data normalization. This left a total of 5953 genes for the final analysis.

To determine whether there was evidence for dosage compensation at the whole-chromosome level, we compared the average FPKM ratio for genes on an aneuploid chromosome to the expectation from changes in gene dose due to changes in chromosome copy number. Thus, a trisomic chromosome would be expected to show a 1.5-fold increase in gene expression and an average FPKM ratio = 1.5 (log2ratio = 0.585). Similarly, monosomic and tetrasomic chromosomes should show average FPKM ratios of 0.5 (log2ratio = -1) and 2 (log2ratio = 1), respectively. We asked whether the observed distribution was consistent with the expected FPKM ratio by calculating the mean and confidence interval of the average FPKM ratio (a one-sample t-test). All analyses were done in RStudio (Team 2013). R scripts for data analysis are available at https://github.com/hollygene/Dosage_Compensation/tree/master/src/R/scripts/Final_Scripts_Used.

As we did for chromosome-level gene expression analysis, previous studies have almost exclusively used FPKM to measure gene expression to compare across strains or treatments. However, the use of FPKMs has been criticized because of loss of power when there are relatively few replicates (three in our experiments), they can vary between samples, and they can be affected by different reference transcriptome annotations (Zheng *et al.* 2011; Wagner *et al.* 2012; Wu *et al.* 2013; Arora *et al.* 2020). The possible loss of statistical power is tied to the multiple comparison issue that arises when examining genes one at a time; however, this is not an issue when comparing chromosome-level expression with tens or hundreds of genes for each chromosome as we did in the chromosome-wide gene expression analysis above. We thus used FPKM data to make our results more comparable to previous studies. When examining individual genes, however, power becomes a very serious concern. As such, we used an alternate method, *DESeq2* (Love *et al.* 2014), that uses statistical models to estimate the distribution for the expression level for a gene in a particular treatment (in our case ancestor versus MA line). Importantly, the method also models the dispersion of the read depth (RNA expression level), assuming the distribution of the expression level can be accurately represented by a negative binomial so that genes of similar expression have similar dispersion. This method is thus expected to more accurately estimate the actual read depth by explicitly considering the variance in the read depth across replicates. As a result, an unusually high or low depth for one replicate will have less impact on the normalization compared to the depths for the other replicates. The method should thus be able to better detect genes that are differentially expressed (DE) in an MA line versus its ancestor when analyzing individual genes.

Raw read counts obtained from Htseq-count were used as input for DESeq2 (Love *et al.* 2014; Anders *et al.* 2015). Individual *DESeqDataSets* were produced for each strain, due to the high variation found across strains, as determined by principal component analysis (PCA) (Supplemental Figure 1). Genes expressed at low levels tend to have high variance and there is thus low power to detect changes in expression. Reads with counts less than 10 in every replicate were removed from further analysis. We used a more stringent cutoff in this analysis to focus on genes for which we have the most power for detecting a change in expression, since we are analyzing individual genes. Removing such genes from the data set resulted in 5532 genes being analyzed.

The *DESeq()* function was implemented on all datasets with default parameters. Annotations were added using the *S. cerevisiae* database from Bioconductor (Carlson M 2015). The *results()* function in *DESeq2* was implemented with default parameters, using a False Discovery Rate (FDR) of 0.1. Analyses were performed with one MA line and the ancestor at a time, since running all strains together would lead to an overestimate of dispersion because of the numerous aneuploid chromosomes in MA lines (see above). For one of the two batches of the homozygous strain, one of the ancestor replicates was substantially different based on a PCA, and so only 2 of the 3 ancestor replicates were used (Supplemental Figure 1A). Similar to the whole chromosome analysis, ratio distributions equal to the sample mean divided by the ancestral mean for the normalized counts were obtained from *DESeq2* estimated read counts. To visualize the data, histograms for both cis (present on aneuploid chromosome) and trans (present on remainder of chromosomes) genes were generated using ggplot2 in R (Wickham 2016).

In addition to looking at all genes in the genome to identify those that were differentially expressed, we also specifically concentrated on a few classes of genes that have been identified in previous work as either being dosage sensitive (DS) (115 genes, Makanae *et al.* 2013), or particularly likely to alter expression in response to stress. These latter genes include those in the environmental stress response (ESR) pathway (139 genes, Gasch *et al.* 2000) and those thought to play a role in aneuploidy stress response (ASR) (201 genes, Torres *et al.* 2007). ASR genes were previously shown to be significantly differentially expressed in aneuploid but not euploid strains. To identify significant DE for genes from these categories, we tested each gene’s expression against the expected expression for a disomic gene in strains where the genes were not on the aneuploid chromosome(s), and determined which genes were significantly different. We then parsed the significantly differentially expressed genes into the ESR/DS/ASR pathways and counted how many times each gene appeared (across samples; i.e. 7/10 aneuploid lines shared 3 DE ASR genes) as a measure of its degree of consistent DE across aneuploid MA lines.

### Data availability

Raw sequencing reads and processed data files used in this analysis are available under the GEO accession number ####### (will be available once manuscript is published).

## Results

### The rate of spontaneous aneuploidy is nearly twice as high in the heterozygous strain as the homozygous strain

The number of aneuploidy events by chromosome is shown in Table 1. We assume that aneuploidy is caused by mitotic nondisjunction since cells were kept asexual. Even if a cell attempted to undergo meiosis, the cessation of growth, coupled with the short time between transfers would essentially guarantee that it would be lost during passaging. In addition, in the heterozygous ancestor we saw no cases of close-to-50% genome-wide homozygosity, which would be expected with meiosis and then intratetrad mating to regenerate a diploid strain (data not shown). The total number of events in the homozygous ancestor strain varied between 0 and 5 per chromosome, which implies a maximal observed rate of nondisjunction for a single chromosome of 1.70 × 10^−5^ events/division (obtained by solving (1-*μ*_*c*_)^2063^ = 140/145 for *μ*_*c*_), and a minimum of zero. The observed rate of an event for any chromosome (i.e. the genome-wide rate) is 6.73 × 10^−6^ events/division (obtained by solving (1-*μ*)^2063^ = 1 – 32/(16*145) for *μ*_*c*_). The total number of events in the heterozygous ancestor varied between 0 and 7 per chromosome, which implies a maximal observed rate of nondisjunction for a single chromosome of 1.56 × 10^−4^ events/division (obtained by solving (1-*μ*_*c*_)^2108^ = 69/76 for *μ*_*c*_), and a minimum of zero. The observed rate of an event for any chromosome is 1.51 × 10^−5^ events/division (obtained by solving (1-*μ*)^2108^ = 1 - 38/(16*76) for *μ*c), which is over two-fold higher than for the homozygous strain. Examination of the number of euploid versus aneuploid lines indicates that this is a significant difference (Fisher’s Exact test, p = 0.004102).

**Table 1.** The number of monosomies, disomies and tetrasomies seen for each MA experiment. The homozygous MA experiment (HH) had 32 events in 145 MA lines maintained for 2063 generations. The heterozygous MA (Hh) experiment had 38 events in 76 MA lines maintained for 2103 generations.

| Chrom. # | # monosomic |  | # trisomic |  | # tetrasomic |  | total events |  |
| --- | --- | --- | --- | --- | --- | --- | --- | --- |
|  | HH | Hh | HH | Hh | HH | Hh | HH | Hh |
| 1 | 0 | 1 | 1 | 3 | 0 | 0 | 1 | 4 |
| 2 | 0 | 0 | 3 | 0 | 0 | 0 | 3 | 0 |
| 3 | 0 | 0 | 2 | 0 | 0 | 0 | 2 | 0 |
| 4 | 0 | 0 | 3 | 0 | 0 | 0 | 3 | 0 |
| 5 | 0 | 0 | 3 | 4 | 0 | 0 | 3 | 4 |
| 6 | 0 | 0 | 0 | 0 | 0 | 0 | 0 | 0 |
| 7 | 0 | 0 | 1 | 5 | 0 | 0 | 1 | 5 |
| 8 | 0 | 0 | 4 | 0 | 0 | 0 | 4 | 0 |
| 9 | 2 | 0 | 3 | 2 | 0 | 0 | 5 | 2 |
| 10 | 0 | 0 | 1 | 1 | 0 | 0 | 1 | 1 |
| 11 | 0 | 0 | 1 | 0 | 0 | 0 | 1 | 0 |
| 12 | 0 | 0 | 1 | 7 | 0 | 0 | 1 | 7 |
| 13 | 0 | 0 | 0 | 0 | 0 | 0 | 0 | 0 |
| 14 | 0 | 0 | 4 | 2 | 0 | 0 | 4 | 2 |
| 15 | 0 | 0 | 0 | 1 | 0 | 0 | 0 | 1 |
| 16 | 0 | 0 | 3 | 10 | 0 | 1 | 3 | 11 |

We note that there were two monosomies and 30 trisomies, a 15-fold difference, in the homozygous experiment and one monosomy and 35 trisomy events in the heterozygous experiment. Since a single nondisjunction event creates both types of aneuploids in the daughter cells, this imbalance implies that monosomies are substantially under-represented in the MA experiments. This finding suggests that monosomies have effects on fitness that are large enough to be seen by selection, even in the low-selection MA framework. Thus, the actual rate of aneuploidy might perhaps be better estimated as twice the trisomy event rate, giving 1.23 × 10^−5^ and 2.77 × 10^−5^ events per cell division for the homozygous and heterozygous ancestor strains, respectively.

In addition, two chromosomes, 6 and 13, comprise 0 out of 70 observed aneuploidy events across the two experiments. If events occurred at random, each chromosome should have 1/16 of the observed events, or 4.4 each. Under a Poisson distribution, the probability of having a chromosome with no events when the expected number is 4.4 equals *e*^−4.4^ = 0.013. It thus seems clear that aneuploidy of chromosomes 6 and 13 either cause strongly deleterious fitness effects or are not tolerated (i.e. are lethal). However, these aneuploidies have been seen in clinical yeast samples (Zhu *et al.* 2016), suggesting that differences in genetic background and/or environment may alter the degree to which aneuploidy is deleterious.

To address whether one aneuploidy event increases the probability of another, we asked whether there was an excess of strains carrying two or more aneuploidies. For the homozygous strain, 28 of the 145 MA lines were found to be aneuploid. Of these, four lines contained two aneuploidies (i.e. two separate chromosomes had become aneuploid), which is not significantly different from the Poisson expectation of 3 (Fisher’s Exact Test, p > 0.99). For the heterozygous strain, 29 out of 76 sequenced MA lines were found to be aneuploid. Of these, seven lines contained two aneuploidies, which is the same as the Poisson expectation. These results suggest that one aneuploidy event does not increase the probability of another. Similarly, we found only one tetrasomic sample across the two experiments. A single non-disjunction event can produce both a monosomic and a trisomic chromosome in a diploid strain. Thus, two events are required to have obtained the one tetrasomic MA line.

To determine whether chromosome size effects the number of nondisjunction events observed in our MA experiments, we plotted size versus number of nondisjunction events (Supplemental Figure 2). While there is clearly variation in which chromosomes become aneuploid, there was no significant relationship with size (R^2^=0.028 heterozygous ancestor; R^2^=0.046 homozygous ancestor). However, only two chromosomes, I and IX, were found to be monosomic and both of these are relatively small chromosomes (I is the smallest at 230,218 bp, and IX is the 4^th^ smallest at 439,888 bp). This suggests that while there is no noticeable effect of chromosome length on aneuploidy occurrence, monosomy may be better tolerated for smaller chromosomes than larger chromosomes.

### Little evidence for whole-chromosome dosage compensation in either strain

We performed RNAseq on 10 euploid and 12 aneuploid homozygous ancestor strain MA lines, and on 6 euploid and 10 aneuploid heterozygous ancestor strain MA lines. Whole-chromosome gene expression was analyzed by calculating the average and 95% confidence intervals of gene expression for each chromosome (Figure 1, Supplemental Figure 4). ANOVAs were also run on each aneuploid sample, comparing the average gene expression from each chromosome to that of the other samples (*lm(y~Line)*, where *y* is FPKM ratio and *Line* is the line number). If there were complete dosage compensation occurring on the whole-chromosome level, we would expect no difference between aneuploid and euploid chromosomes, such that ANOVAs would show no effect of chromosome number on gene expression. However, in the absence of dosage compensation, chromosome number would have an effect, with aneuploid chromosomes underlying the significant difference among chromosomes. Further, in the absence of dosage compensation, we would expect gene expression to mirror gene dose such that aneuploid chromosomes would show 0.5 or 1.5-fold increases in expression for monosomic and trisomic chromosomes, respectively.

**Figure 1:**
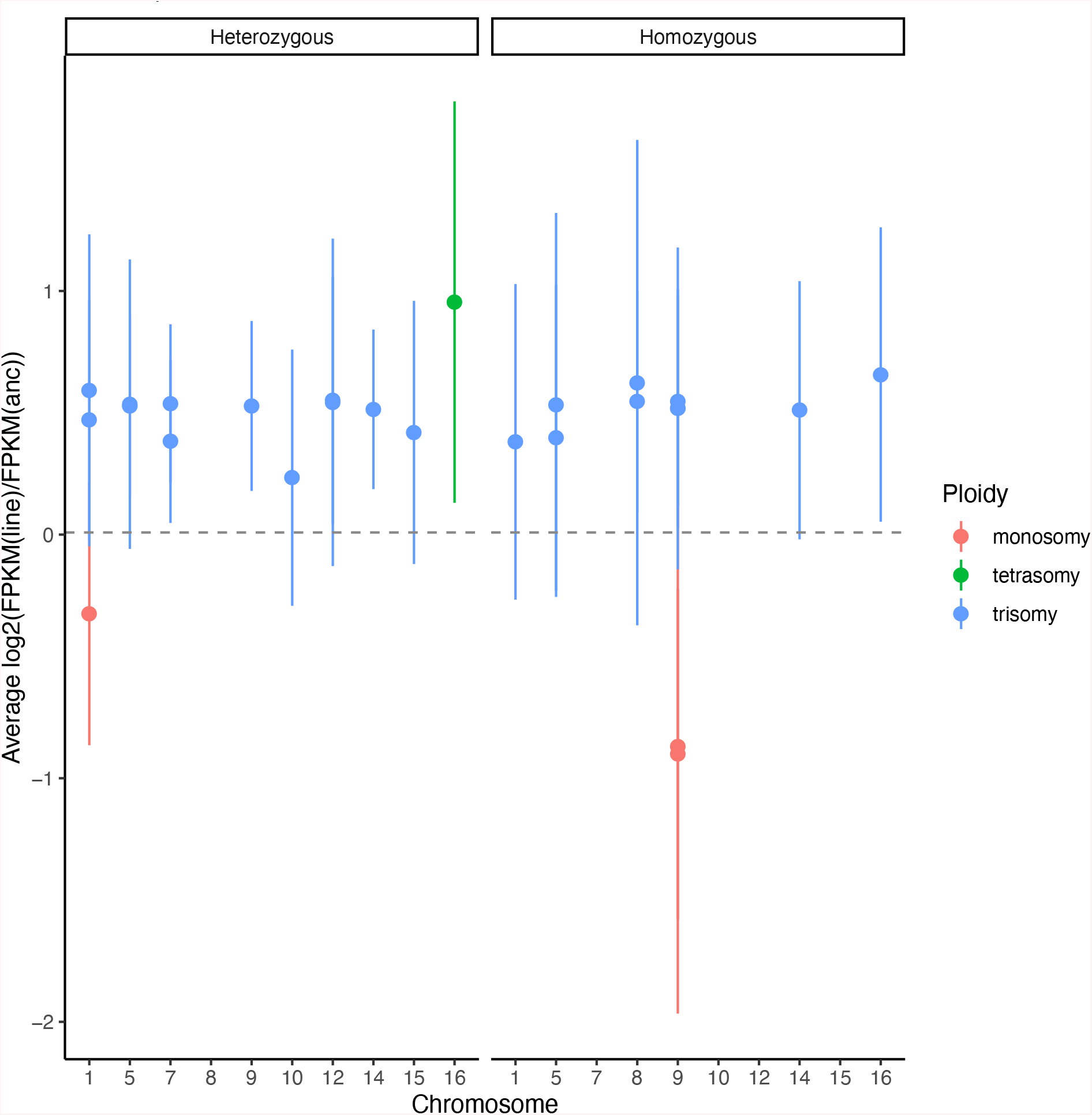
Average log2 ratio of FPKM values in each aneuploid line compared to their respective ancestor. Dashed gray line is the average log2 ratio of FPKM values of all the euploid lines combined. Error bars are +/− one standard deviation.

ANOVAs indicated that the effect of chromosome was significant (p < 0.01) in every aneuploid MA line, as expected with no dosage compensation (Supplemental Data). For chromosomes that did not have any aneuploid lines represented in the dataset, we still found some differential expression in a few aneuploid lines. Specifically, for chromosome III, Line 76, 61, 59, 49, 18, 11 (heterozygous ancestor) are significantly different (*p*<0.01), suggesting that aneuploidy causes changes in gene expression of genes on chromosome III across aneuploid strains. Of the genes on chromosome III, 3 are in the aneuploid stress response (YCL037C, YCR057C, and YCR072C), 4 are in the environmental stress response (YCL035C, YCL040W, YCR004C, YCR091W), and 1 is dosage sensitive (YCR088W). However, none of the environmental or aneuploid stress response genes on this chromosome were significant in more than one sample from either ancestor, implying that the aneuploid/environmental stress response is unique to each sample. This might be explained by the samples having different aneuploidies – only samples 59 and 61 shared the same trisomy.

ANOVAs on some euploid lines also gave significant *p* values for certain chromosomes, indicating that some chromosomes show changes in expression even in the absence of aneuploidy (Figure 1, Supplemental Figure 4, Supplemental Data). This could suggest an impact of passaging yeast in an MA framework on gene expression in yeast, but it is important to note that this result is from FPKM data, which can vary greatly between samples (see above).

If there were no chromosome-level dosage compensation, then the level of gene expression is expected to be proportional to chromosome copy number. For most aneuploid chromosomes in MA lines this prediction held: expression levels did not differ significantly from the expectation. However, in 4 MA lines (line numbers 18, 49, 59 and 61) from the heterozygous ancestor, the expected level of gene expression was less extreme than expected based on chromosome copy number (Figure 1, Supplemental Figure 4). Chromosome I of line 18 had average expression change equal to 1.3-fold, chromosome V of line 49 had average expression change equal to 1.35-fold, chromosome VII of line 59 had average expression change equal to 1.25-fold, and chromosome VII of line 61 had average expression change equal to 1.39-fold. All these values were significantly different from the expected expression level of 1.5-fold (*p* < 0.05, one-sample t-test). The vast majority of gene expression changes, 65 of 69 aneuploid chromosomes, are consistent with a lack of whole-chromosome dosage compensation occurring in either strain, and together these findings support previous work showing no whole-chromosome dosage compensation in aneuploid yeast (Torres *et al.* 2010).

### Distribution of gene expression from euploid versus aneuploid chromosomes

The previous analysis indicates that mean gene expression of aneuploid chromosomes seems to be predicted by gene dose. We next examined whether the mean expression for genes on the non-aneuploid (disomic) chromosomes is altered by aneuploidy. In addition, we examined whether the variance in gene expression for aneuploid chromosomes is the same as for euploid chromosomes in the 20 aneuploid MA lines used for RNA sequencing, and whether the variance in gene expression differs between euploid MA lines and their euploid ancestor. The distribution of FPKM ratios (MA line FPKM / ancestor FPKM) for all genes in euploid samples (Supplemental Figures 5 & 6), for genes on the aneuploid chromosome(s) in aneuploid samples (cis genes), and for genes not located on the aneuploid chromosome(s) in aneuploid samples (trans genes) were analyzed (Supplemental Figures 7-12).

The expected mean expression ratio in euploid lines is 1. In every euploid line analyzed, the expected distribution had a mean that was indistinguishable from 1 (*p*> 0.1, Supplemental Figures 5 & 6). For aneuploid lines, the expected mean expression for trans genes (those not located on the aneuploid chromosome) is not equal to 1. This is because the aneuploid chromosome will have more (for trisomy) or fewer (for monosomy) reads mapping to it than in a euploid line. This changes the percentage of reads that map to other chromosomes – fewer for trisomy and more for monosomy. The extent of this effect will also depend on chromosome size because chromosome size alters the percentage of the genome that it represents, with aneuploidy for larger chromosomes having a larger effect on the reads mapping to other chromosomes. In Supplemental Table 1, we indicate the expected mean lower expression level for trans genes in MA lines carrying a single trisomy. Similarly, for lines with monosomies, the expected higher mean expression of trans genes is indicated. We tested the mean expression of trans genes against the expectation based on the chromosomes for which they were aneuploid and found that in no case were they significantly different (*p* > 0.05, one-sample t-test; Supplemental Figures 7-12).

To examine whether the variance in gene expression is greater in aneuploid lines, we compared the variance in gene expression of both cis and trans genes to the variance of those same genes in a euploid line using a Levene’s test. For comparisons, we randomly matched a euploid line with each aneuploid line. We determined whether the means and variances of these distributions differed from the expectation (the expectation being that both the means and the variances are equal between aneuploid and euploid lines). The variances of gene expression of cis genes were significantly different from the expectation in every case except for two: the comparison of homozygous line 15 (trisomic for chromosome 9) to homozygous line 5 (euploid) and the comparison of homozygous line 152 (trisomic for chromosomes 1 and 7) to homozygous line 1 (euploid) (Supplemental Table 2). There is nothing immediately notable with these samples, though the ANOVA for chromosome VII line 1 was significant (*p*<0.05); there was no similar connection in line 5 for chromosome IX (Supplemental Data).

### Individual Dosage-Compensated Genes

Our analyses indicated that at the whole-chromosome level aneuploidy leads to changes in gene expression predicted by gene dose, such that there was no evidence for dosage compensation, and minor (or no) effects on expression of the rest of genome. Next, we investigated individual genes. We sought to group genes present on aneuploid chromosomes into five categories based on their gene expression, using similar metrics as a previous study (Malone *et al.* 2012): 1. Not dosage compensated: these genes have expression levels not significantly different from those predicted by their gene dose. 2. Partially dosage compensated: these genes show less-extreme gene expression changes than predicted by their dose. 3. Fully dosage compensated: these genes show no change in expression in response to changes in gene dose. 4. Over-dosage compensated: these genes show changes in expression that are in the opposite direction of the change in gene dose. 5. Anti-dosage compensated genes show more extreme changes in expression than predicted by the change in gene dose (in the direction of the aneuploidy – i.e. monosomic genes would have lower gene expression than predicted by monosomy) (Supplemental Table 3). Any gene that had expression levels different from the ancestor and different from the expectation based on gene dose was assigned to one of the categories depending on their level of expression. For the aneuploid strains we analyzed, we found several genes in each of these categories (Table 2). Since we are testing many genes (5532), power becomes limited due to the need to correct for multiple testing. For this reason, it is important to test for expression that is consistent both with respect to the ancestor and to the expectation based on gene dose. Many genes do not differ from either, in which case we cannot conclude the degree to which they are compensated – these genes were assigned as category 0 genes, or “unknown” compensation. Our analyses revealed that the power to distinguish whether a gene exhibits dosage compensation or not is low; the vast majority of genes are in category 0 (Table 2). For those genes in other categories, we find that there is little agreement between different strains in terms of the percentage of genes in these categories.

**Table 2.** The number of genes in each expression change category across the aneuploid strains for which we have RNAseq data. 0 = unknown, 1 = no dosage compensation (DC), 2 = partial DC, 3 = full DC, 4 = over-compensation, 5 = anti-compensation. Key in supplement. HH: Homozygous ancestor; Hh: heterozygous ancestor

| MA line | Aneuploidy | Category |  |  |  |  |  |
| --- | --- | --- | --- | --- | --- | --- | --- |
|  |  | 0 | 1 | 2 | 3 | 4 | 5 |
| **HH-152** | 1, 7 |  |  |  |  |  |  |
|  | Chr 1: | 52 | 22 | 6 | 0 | 0 | 1 |
|  | Chr 7: | 482 | 9 | 12 | 0 | 0 | 0 |
| **HH-117** | 5 | 147 | 107 | 5 | 0 | 0 | 0 |
| **HH-123** | 5 | 209 | 50 | 5 | 0 | 0 | 1 |
| **HH-108** | 8, 9<sup>m</sup> |  |  |  |  |  |  |
|  | Chr 8: | 212 | 76 | 129 | 0 | 39 | 42 |
|  | Chr 9: | 41 | 8 | 64 | 0 | 84 | 2 |
| **HH-15** | 9 | 62 | 82 | 52 | 0 | 0 | 3 |
| **HH-29** | 9<sup>m</sup> | 90 | 1 | 107 | 0 | 1 | 0 |
| **HH-88** | 9 | 58 | 78 | 50 | 0 | 0 | 13 |
| **HH-119** | 9 | 70 | 92 | 37 | 0 | 0 | 1 |
| **HH-9** | 14 | 92 | 140 | 128 | 0 | 0 | 13 |
| **HH-112** | 16 | 244 | 20 | 0 | 0 | 0 | 0 |
| **Hh-7** | 1 | 51 | 29 | 1 | 0 | 0 | 0 |
| **Hh-11** | 1<sup>m</sup>, 15 |  |  |  |  |  |  |
|  | Chr 1: | 67 | 0 | 10 | 0 | 4 | 0 |
|  | Chr 15: | 413 | 57 | 5 | 0 | 0 | 0 |
| **Hh-18** | 1 | 45 | 29 | 6 | 0 | 1 | 0 |
|  | 12 | 240 | 153 | 42 | 0 | 1 | 26 |
| **Hh-4** | 5 | 165 | 71 | 5 | 0 | 3 | 21 |
| **Hh-49** | 5 | 135 | 89 | 10 | 0 | 0 | 0 |
| **Hh-59** | 7 | 288 | 167 | 23 | 0 | 2 | 13 |
| **Hh-61** | 7 | 160 | 193 | 20 | 0 | 2 | 18 |
| **Hh-76** | 9, 10, 14<sup>p</sup> |  |  |  |  |  |  |
|  | Chr 9: | 76 | 77 | 43 | 0 | 0 | 3 |
|  | Chr 10: | 250 | 62 | 20 | 0 | 0 | 5 |
|  | Chr 14: | 165 | 132 | 81 | 0 | 0 | 2 |
| **Hh-77** | 12 | 145 | 184 | 125 | 0 | 0 | 11 |
| **Hh-8** | 16 | 117 | 126 | 31 | 0 | 16 | 156 |

We compared the trans genes of aneuploid samples with those of samples with a different aneuploid chromosome(s) to determine if there was a common response to aneuploidy, as has been shown in previous studies (Gasch *et al.* 2000; Zillikens *et al.* 2017a). We found that in lines from the heterozygous ancestor, at most, 8/10 aneuploid samples shared 15 of the same DE trans genes (genes that were not located on an aneuploid chromosome). Of these, 2 were in the environmental stress response (ESR) (YIR038C and YKR076W), and one was in the aneuploidy stress response (ASR) (YBR117C) (Gasch *et al.* 2000; Torres *et al.* 2010). In lines from the homozygous ancestor, at most 6/10 aneuploid lines shared 8 of the same DE trans genes. None of these genes were in either the ESR or the ASR.

We then examined if euploid lines shared a common gene expression response and found that in lines from the homozygous ancestor, at most 5/10 euploid samples shared 8 common differentially expressed genes. Of these, one is in the ASR (YOL126C), and none were in the ESR. In the heterozygous ancestor, at most 5/6 lines shared 54 DE genes. Of these, 9 were in the ESR (YBR026C, YCR004C, YFR053C, YGR043C, YKL026C, YKR009C, YLL026W, YMR110C, and YOR374W). These genes are implicated in metabolic processes, according to GO analysis (Supplemental Figure 13). This result suggests a shared effect of the mutation accumulation experimental design on gene expression, particularly in the heterozygous ancestor lines, with metabolism being the most impacted process.

### Histone Genes

Histone genes H2A and H2B are known to possess a mechanism of dosage compensation in *S. cerevisiae* (Osley and Hereford 1981; Medici *et al.* 2014). Our analyses did not include samples with aneuploidies on the chromosomes containing H2A and H2B (II and IV), but we do have aneuploid samples with RNA sequencing for chromosomes containing other histone genes: XIV, XV, and XVI (containing histones 3,4, and linker, respectively). Six lines across both experiments are trisomic for chromosome XIV, 1 line is trisomic for chromosome XV, 13 lines are trisomic for XVI, and 1 line is tetrasomic for XVI. Previous studies have found that these genes do not display dosage compensation and we also did not find evidence for compensation (Peter R. Eriksson 2012) (Supplemental Table 4).

### Stress Response Genes

Yeast are known to undergo what is known as the environmental stress response (Gasch *et al.* 2000; Zillikens *et al.* 2017a), when conditions are unfavorable due to various factors, including temperature stress, oxidative stress, and nutrient limitation. We analyzed genes previously found to relate to the environmental stress response and found that our aneuploid samples did differentially express most of these genes (Figure 3), though there was no significant trend of a shared differential expression response of ESR genes between samples.

It has been found that similarly, aneuploid yeast undergo what is referred to as the “aneuploid stress response (ASR),” in which certain trans genes are differentially expressed (Torres *et al.* 2010). A majority of these genes are also differentially expressed during the environmental stress response. To determine if we found the same pattern of differential expression in our spontaneously aneuploid samples, we investigated these ASR genes (201 genes total) and found that in samples from the heterozygous ancestor, at most 7/10 aneuploid lines shared 3 DE ASR genes. In the homozygous ancestor aneuploid lines, at most only 4/10 aneuploid lines shared just 1 DE ASR gene. As expected, the euploid lines in both datasets did not show significant signatures of differential expression on ASR genes and as expected, did not share many DE ASR genes (Figure 3).

### Dosage-Sensitive Genes

Previous studies have found that certain genes are more sensitive to changes in gene dose than others. Using the “genetic tug-of-war” method, Makanae et al 2013 found the copy-number limits of overexpression in all 5806 protein-coding genes in *S. cerevisiae*, and found 115 genes whose copy number limits were 10 or less (more than this amount caused cell death) (Makanae *et al.* 2013). Curious as to whether our samples exhibited a compensatory response for these dosage sensitive genes, we looked at the same set of genes and parsed out those that were significantly differentially expressed in our aneuploid samples. Most aneuploid samples had few differentially expressed dosage sensitive genes (Figure 3). The euploid lines in both experiments had very few DE dosage-sensitive genes, consistent with the expectation that there would be zero (Figure 3).

Of particular interest were the genes on the aneuploid chromosomes, as they differ in copy number compared to the rest of the genes in the genome. Most samples showed a high level of compensation of dosage-sensitive genes on the aneuploid chromosome and elsewhere in the genome. However, samples with a trisomy for chromosome 9 appeared to be more tolerant of the duplication (likely due to individual gene compensation) than other chromosomes – samples ranged from 0 to 33% compensation (Table 2).

## Discussion

### Rate of aneuploidy

We calculated the rate of aneuploidy based on data from two previous yeast mutation accumulation experiments passaged for similar numbers of cell generations: one with a heterozygous strain and one with a homozygous strain. We found that the rate of aneuploidy is higher in the heterozygous strain than the homozygous strain (p<0.0001, Fisher’s exact test). The heterozygous ancestor strain MA lines had a total of 29 aneuploids and 47 euploids, whereas the homozygous ancestor MA lines had a total of 28 aneuploid and 117 euploid lines. Previous studies have found that hybrids of two yeast species systematically lose all or part of one parent’s genome (Marinoni *et al.* 1999). Since the heterozygous ancestor strain had a heterozygous site every ~250 bases, it is possible that the mating of distantly related *S. cerevisiae* strains to produce the heterozygous strain showed a milder version of genome incompatibility as exemplified by the higher rate of aneuploidy compared to the homozygous lab strain. However, the heterozygous strain did not show any growth defects (which could have indicated a phenotypic effect of genome incompatibility) compared to the homozygous strain (data not shown). In addition, the homozygous strain ancestor carried an *ade2* mutation, which was found in a recent study to lower a strain’s tolerance to aneuploidy (Hose *et al.* 2020). It is therefore virtually impossible to distinguish the influence of heterozygosity from the influence of this mutation on aneuploidy rate in our strains – the genetic differences between the two are too great and have too much of an influence on aneuploidy tolerance. However, recent RNA sequencing studies of wild yeast have found that heterozygosity is correlated with aneuploidy (unpublished data). To examine the effect of heterozygosity per se in future experiments, homozygous diploids could be generated from each of the parent strains used to make the heterozygous strain and then used in mutation accumulation experiments to determine the rate at which aneuploidies arise.

In our experiment, we found 3 and 6 events for the homozygous and heterozygous strains involving chromosome V nondisjunction, implying a rate of 9.67 × 10^−6^ and 3.90 × 10^−5^ events per cell division, respectively. Previous studies have found that chromosome V is lost spontaneously by nondisjunction in *S. cerevisiae* at a rate of 2-8 × 10^−6^ cell generations (Mulla *et al.* 2014). This estimate is significantly different from the homozygous strain (*p* = 0.0001) and much more significantly different from the heterozygous strain (*p* < 0.00001). This previous study used a laboratory strain (A364A), which is highly homozygous and has *ade1* and *ade2* auxotrophies, which perhaps explains the discrepancy in rates between their estimate and our heterozygous ancestor estimate.

We found a difference in aneuploidy rates between the ancestor strains at the individual chromosome level as well as overall. In the heterozygous ancestor strain MA lines, we found 10 (out of 29 total aneuploidies) trisomies of chromosome XVI, compared with 3 (out of 29 total aneuploidies) in the homozygous ancestor MA lines (Table 1). Previous studies have found a similar discrepancy between diploid and diploid-hybrid strains of yeast, with the hybrid strains showing a higher rate of aneuploidy at chromosome XVI (Kumaran *et al.* 2013). These results suggest that heterozygosity influences either nondisjunction rate or tolerance of certain aneuploidies and that certain chromosomes are either more likely to become aneuploid or are better tolerated after becoming aneuploid, or both.

Due to the diploid nature of our initial MA ancestors, we were able to analyze trisomics, monosomics, and a tetrasomic to study the rate and effects of whole-chromosome aneuploidy. Contrary to most previous studies, we were able to observe the spontaneous rate and effects of monosomy, which is substantially less common than trisomy in our samples (Table 1). Considering nondisjunction events result in the production of both a trisomy and a monosomy, we would expect to see an equal number of each in our data. The lack of monosomies implies that there must be strong selection against them, implying that fewer copies of a chromosome is substantially more deleterious than additional copies in order to be lost in the MA experimental framework. One explanation could be that a monosomy has a larger effect on gene expression, a two-fold difference, compared to trisomy, which results in a 1.5-fold difference. Tetrasomies are also a 2-fold difference but require two events, which may explain their rareness.

### No evidence for whole-chromosome dosage compensation at the transcript level

Our results suggest that there is no general mechanism for dosage-compensation in aneuploid yeast, either at the whole-chromosome or individual gene level (Supplemental Figure 4,Table 2). Our results mirror previous findings that RNA level scales with DNA copy number and that no RNA-level compensation occurs (Torres *et al.* 2010). This is in contrast to studies that have reported whole-chromosome dosage compensation in yeast (Hose *et al.* 2015; Gasch *et al.* 2016). One explanation that has been proposed for the differences in results is heterogeneous samples containing both aneuploid and euploid cells (Hose *et al.* 2015; Gasch *et al.* 2016), causing gene expression ratios to be intermediate between what is expected for aneuploid and euploid DNA copy levels. A recent study mapped the genetic basis of aneuploidy tolerance in wild yeast to SSD1, an RNA-binding protein involved in proteolysis (Hose *et al.* 2020), suggesting that any compensatory mechanism in wild aneuploid yeast is active at the protein level. The strains used in this study both have fully functional SSD1 proteins, according to translation using Geneious (Supplemental Figure 3), suggesting that the differences in aneuploidy rate are likely not caused by lack of or presence of a functional SSD1 protein. In addition, the apparent partial compensation we observed for some MA strains in our study may be caused by heterogenous samples. However, we feel this is unlikely as a constantly heterogeneous population would likely revert to euploidy – even in the MA framework – as aneuploidy typically causes fitness defects. To avoid this potential problem, future studies could employ the use of fluorescence activated cell sorting (FACS) to separate the aneuploid cells from the euploid cells and use only the aneuploid culture for RNA extraction. Previous studies have found that the increase in a partner gene can rescue the sensitivity of a strain to another with increased dosage. This may be occurring in the samples that had little to no compensation of the dosage sensitive genes on the aneuploid chromosome (Figure 3). Further investigation into these partner genes is required for future studies to determine if this is the case.

### Categorization of Individual Gene Expression

We investigated the effects of aneuploidy on gene expression at the individual-gene level and found that because of low power, it was challenging to detect statistically significant changes in gene expression level for the majority of genes analyzed (see Results), therefore the majority of genes fell into the “unknown” category of compensation (Supplemental Table 3). However, for the genes that have sufficient statistical power, we found that most were not dosage compensated, several were partially compensated, and very few were over- or anti-compensated (Table 2). Future studies will benefit from using more replicates to increase power for individual gene analysis in order to further characterize the large number of genes falling into the “unknown” category.

### Aneuploidy effects on trans genes

Previous studies have proposed that there is an effect of aneuploidy on the remainder of the genome, by looking at the peaks of the distributions and claiming that the apparent skew to the left of 1.00 indicated that the aneuploid chromosome was causing other expression effects in the genome (Hou *et al.* 2018). We investigated trans genes in our data and found that they showed the expected level of gene expression (Figure 2, Supplemental Figures 7-12); trisomies lead to an apparent reduction in expression of trans genes and monosomies lead to an apparent increase, but the shift is as predicted based on the size of the genomes of the aneuploid chromosome implying that aneuploidy does not cause a global change in gene expression. We also determined whether aneuploid lines shared any differentially expressed genes not located on aneuploid chromosomes. We compared gene expression data between aneuploid samples and found, in our heterozygous ancestor, only 15 commonly differentially expressed trans genes among 8 of the 10 aneuploid lines. Of these, 2 were in the environmental stress response (ESR) genes, and one was in the aneuploidy stress response (ASR) genes (Gasch *et al.* 2000; Torres *et al.* 2010). Similarly, in the homozygous ancestor lines, we found only 8 commonly differentially expressed trans genes among 6 of the 10 aneuploid lines. However, none of the differentially expressed trans genes in the homozygous ancestor had been previously identified as sensitive to aneuploid or environmental stress. The discrepancy in differentially expressed genes between the heterozygous and homozygous ancestor samples may be due to different genetic backgrounds or could be impacted by heterozygosity. However, there is no evidence in our study for common transcriptional responses to aneuploidy in spontaneously-aneuploid yeast across genotypes/genetic backgrounds, suggesting that adaptation to aneuploidy is not facilitated by a common compensation mechanism or stress response.

**Figure 2.**
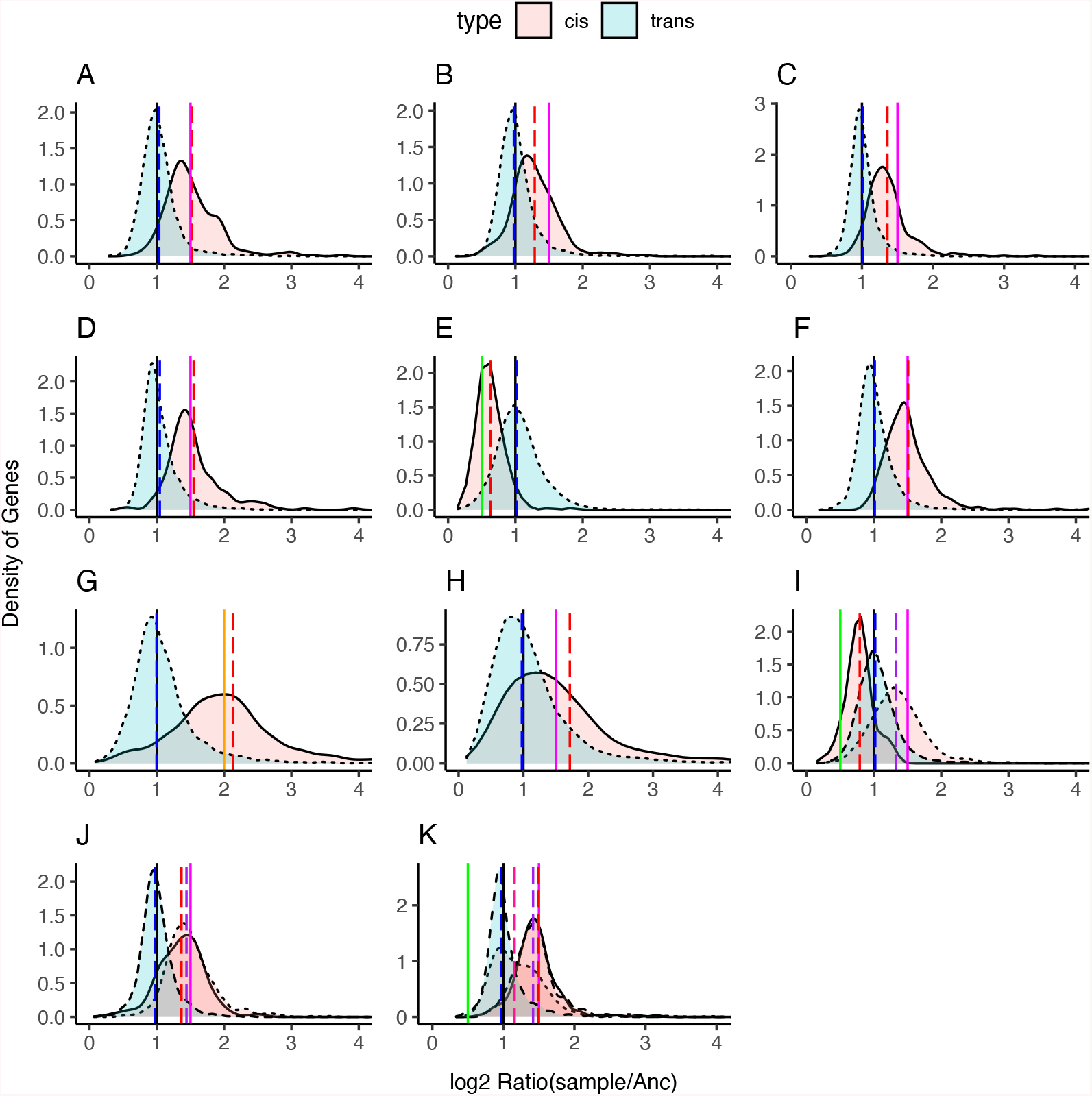
Read count ratio distributions of select lines. Genes on (cis) and off (trans) of the aneuploid chromosome(s) are labeled in pink and blue, respectively. Multiple aneuploid chromosomes are indicated by line type (I,J,K). Vertical lines: red dotted line: mean ratio of cis genes, blue dotted line: mean ratio of trans genes, black line: ratio of 1 (equal expression compared to ancestor), green line: ratio of 0.5 (expectation for monosomic genes), magenta line: ratio of 1.5 (expectation for trisomic genes), orange line: ratio of 2 (expectation for tetrasomic genes). A: Heterozygous ancestor line 4, trisomic for chromosome V; B: homozygous ancestor line 117, trisomic for chromosome V, C: heterozygous ancestor line 59, trisomic for chromosome VII, D: homozygous ancestor line 15, trisomic for chromosome IX; E: homozygous ancestor line 29, monosomic for chromosome IX, F: homozygous ancestor line 9, trisomic for chromosome XIV; G: heterozygous ancestor line 8, tetrasomic for chromosome XVI; H: homozygous ancestor 112, trisomic for XVI; I: Heterozygous ancestor line 11, trisomic for chromosome XV and monosomic for chromosome I; J: heterozygous ancestor line 18, trisomic for chromosome I and XII; K: heterozygous ancestor line 76, trisomic for chromosome IX and XIV, partial duplication of chromosome X (pink dotted line is mean ratio for chromosome X in this line. Larger figures and remaining aneuploid lines are available in the supplement.

**Figure 3:**
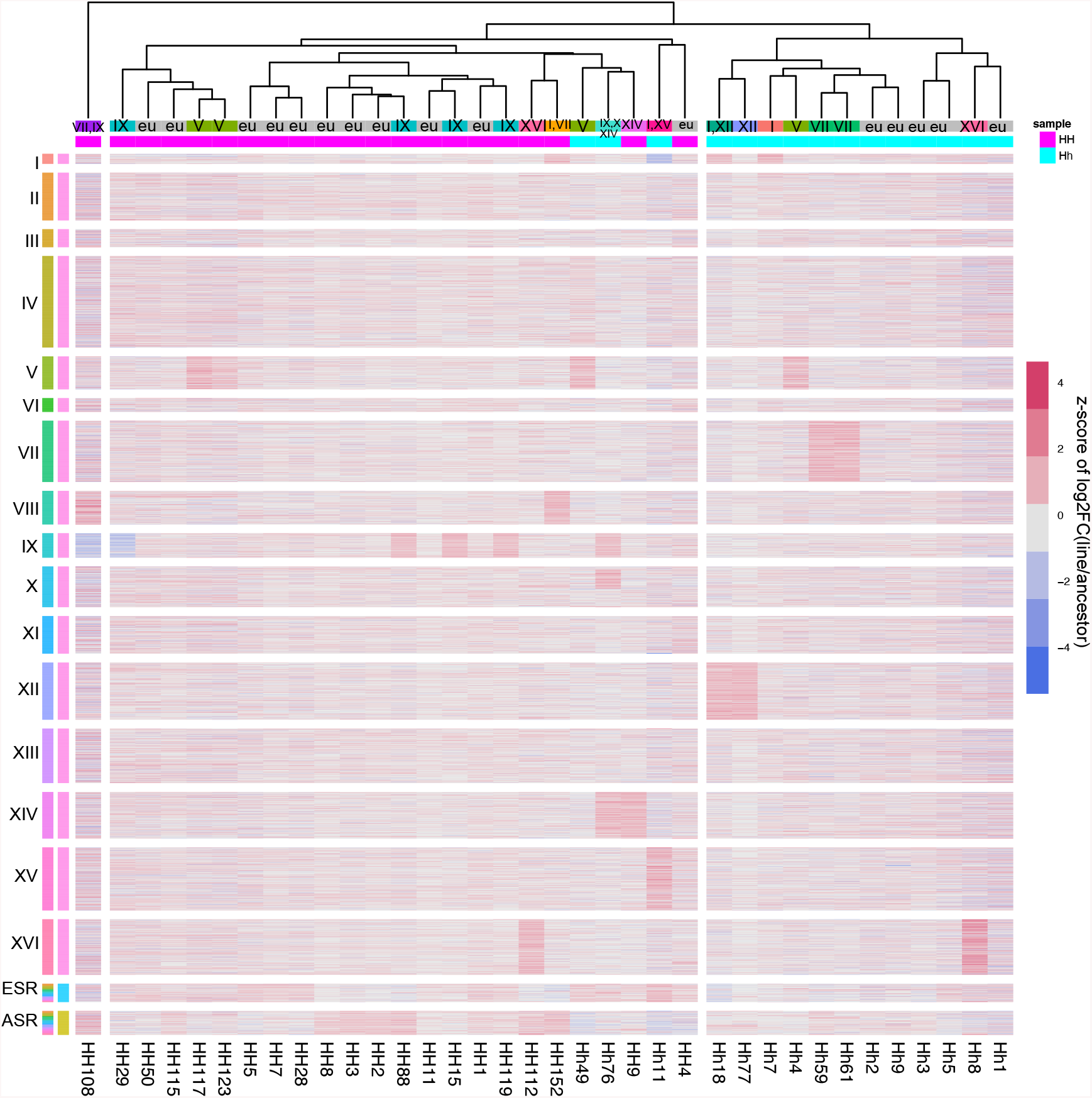
Heatmap depicting all lines sequenced. Each horizontal line represents a single transcript values are z-scores of log2 fold changes of the expression of the line divided by the expression of the ancestor. Individual chromosomes are grouped together, and individual lines are clustered based on similarities in gene expression. Each aneuploid chromosome can be seen as a block of red or blue (red for higher expression and blue for lower expression); it is notable that chromosome X of line 76 shows the half of the chromosome that is duplicated. The ESR and ASR genes are grouped together at the bottom of the heatmap; samples that share differentially expressed genes in these categories can be seen as faintly higher (or lower) gene expression levels.

Previous studies in yeast have found evidence of a transcriptional response to environmental stress as well as a transcriptional response to aneuploidy involving the “environmental stress response” genes and the “aneuploidy stress response” genes, respectively (Gasch *et al.* 2000; Torres *et al.* 2007; Zillikens *et al.* 2017b). We investigated the environmental stress response (ESR) genes and found that most ESR genes were differentially expressed in our aneuploid samples, but not in euploid samples, suggesting that the state of aneuploidy has similar effects on the transcriptome to various environmental stresses including high salinity, high temperatures, and highly oxidative-species rich environments. It would be interesting to know if the yeast samples exposed to these environmental stresses had any copy number changes in their genomes – this would add evidence to the hypothesis that aneuploidy is an adaptive state to changes in the environment and/or a consequence of stress. However, our aneuploid strains do not have many shared differentially expressed aneuploidy stress response genes, suggesting that each aneuploidy confers a different stress and therefore a different transcriptional stress response. This is in contrast to previous studies which found a common transcriptional response to aneuploidy (Torres *et al.* 2007). One explanation for this discrepancy is the way in which aneuploids were generated: the Torres et al study used a chromosome transfer strategy to select for aneuploids after abortive mating, whereas in this study we used samples with aneuploidies that spontaneously arose during mutation accumulation, without a strong force of selection. It is possible that selecting for aneuploids using abortive mating caused the differential expression signature, not the aneuploidy itself.

## Conclusion

This study demonstrated that heterozygosity and *ade2* auxotrophy is correlated with a higher aneuploidy rate, that there is no evidence for whole-chromosome dosage compensation at the transcriptome level in aneuploid yeast, and that aneuploid chromosomes do not significantly influence the gene expression patterns among the rest of the transcriptome. We did find evidence for compensation at the individual gene level for genes that are particularly toxic in high copy numbers, suggesting that cells are able to employ transcriptional compensatory mechanisms to tolerate aneuploidy at least at the individual gene level. Further, our analyses demonstrated evidence for individual aneuploid lines to differentially express environmental and aneuploidy stress response genes. There were not many shared differentially expressed ESR/ASR genes among aneuploid lines, however, implying that each aneuploid line deals with its aneuploidy in a unique manner. Our finding of no global effects of aneuploidy on gene expression is in direct opposition to a recent paper claiming this – however, we showed that the apparent skew of trans genes is actually due to sequencing bias from reads mapping to more (or less) copies of the aneuploid chromosome(s).

Our results bring insights into the effects of aneuploidy on gene expression in budding yeast and can be applied to other species as well. Further, our findings perhaps provide insight into the evolution of sex chromosomes and dosage compensation – the absence of dosage compensation in a single-celled species that exhibits aneuploidy reasonably frequently in nature suggests partial or complete chromosome aneuploidy during sex chromosome evolution may be reasonably well tolerated, even in the absence of a pre-existing dosage-compensating mechanism.

More insights into how wild yeast tolerate aneuploidy are required. A recent study found that the SSD1 gene in yeast is associated with aneuploidy tolerance in wild strains versus lab strains (Hose *et al.* 2020). This gene is a translational repressor and is functional in wild yeast isolates but not in laboratory strains. This implies that wild aneuploid yeast strains can tolerate aneuploidy by attenuating translation of the duplicated genes. This reflected previous work in aneuploid yeast that showed compensation at the protein, but not RNA, level (Noah dephoure 2014). The strains used in our study both have a functional copy of SSD1, which could explain the relatively high tolerance of aneuploidies in both strains. Our analyses provide further evidence for the lack of transcript-level dosage compensation, and future studies could use SSD1 knockout strains of yeast for mutation accumulation studies and determine rates and tolerance of aneuploidy in a similar manner as this study.

## Supporting information

ANOVAs

