## Supplemental Material for "No evidence for whole-chromosome dosage compensation or global transcriptomic expression differences in spontaneously-aneuploid mutation accumulation lines of *Saccharomyces cerevisiae*"

**Supplemental Figure 1:** Principal components analysis of all lines sequenced.

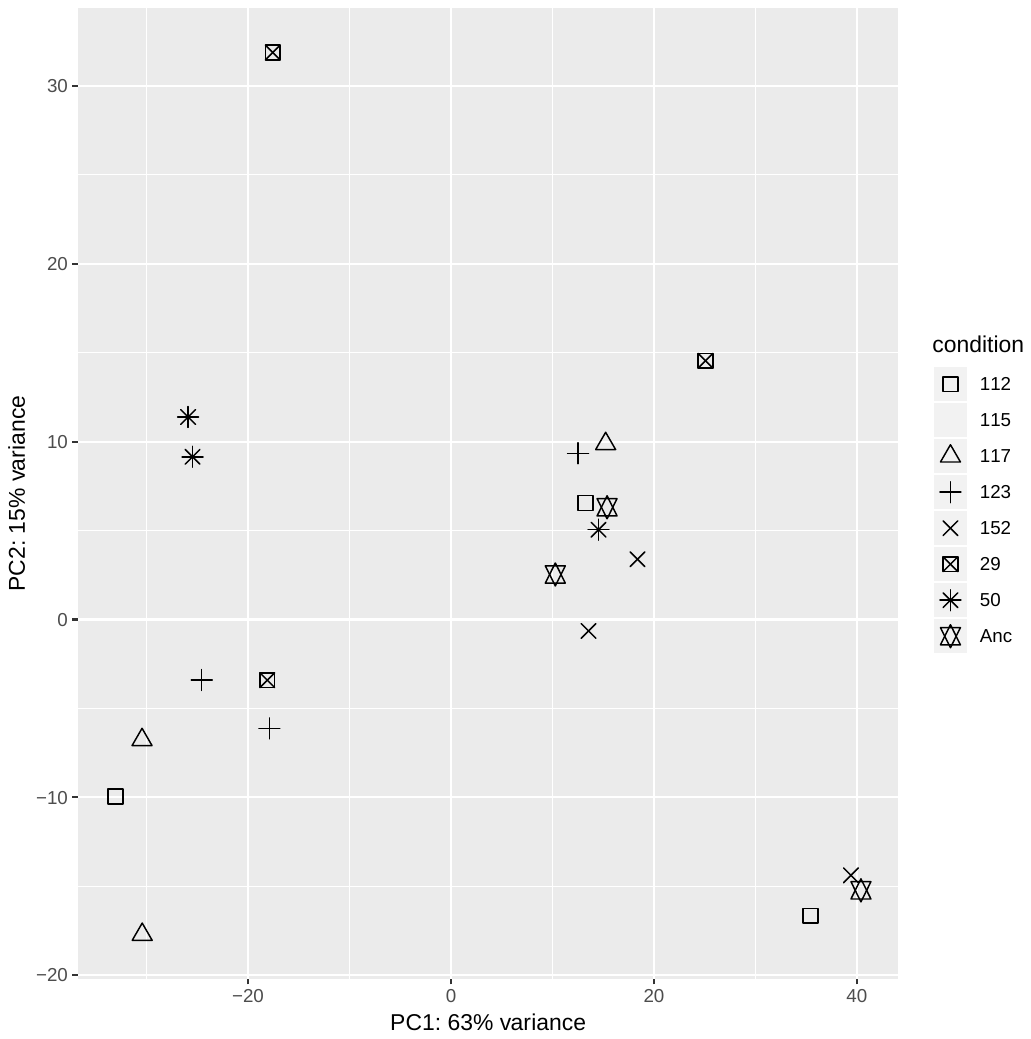

A: PCA of second batch of homozygous ancestor samples. One ancestral replicate (near bottom right) was removed from analysis based on PC1.

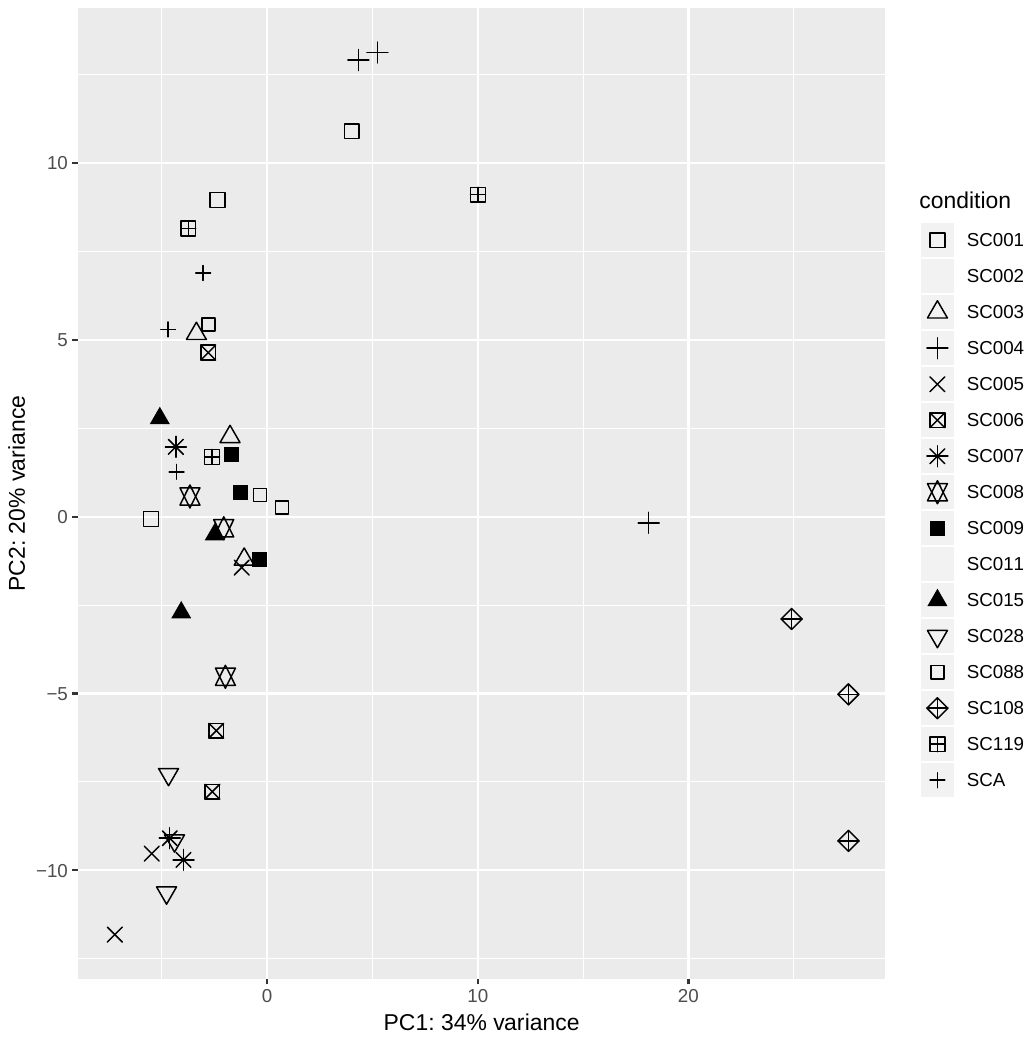

B: PCA of first homozygous ancestor sequencing run.

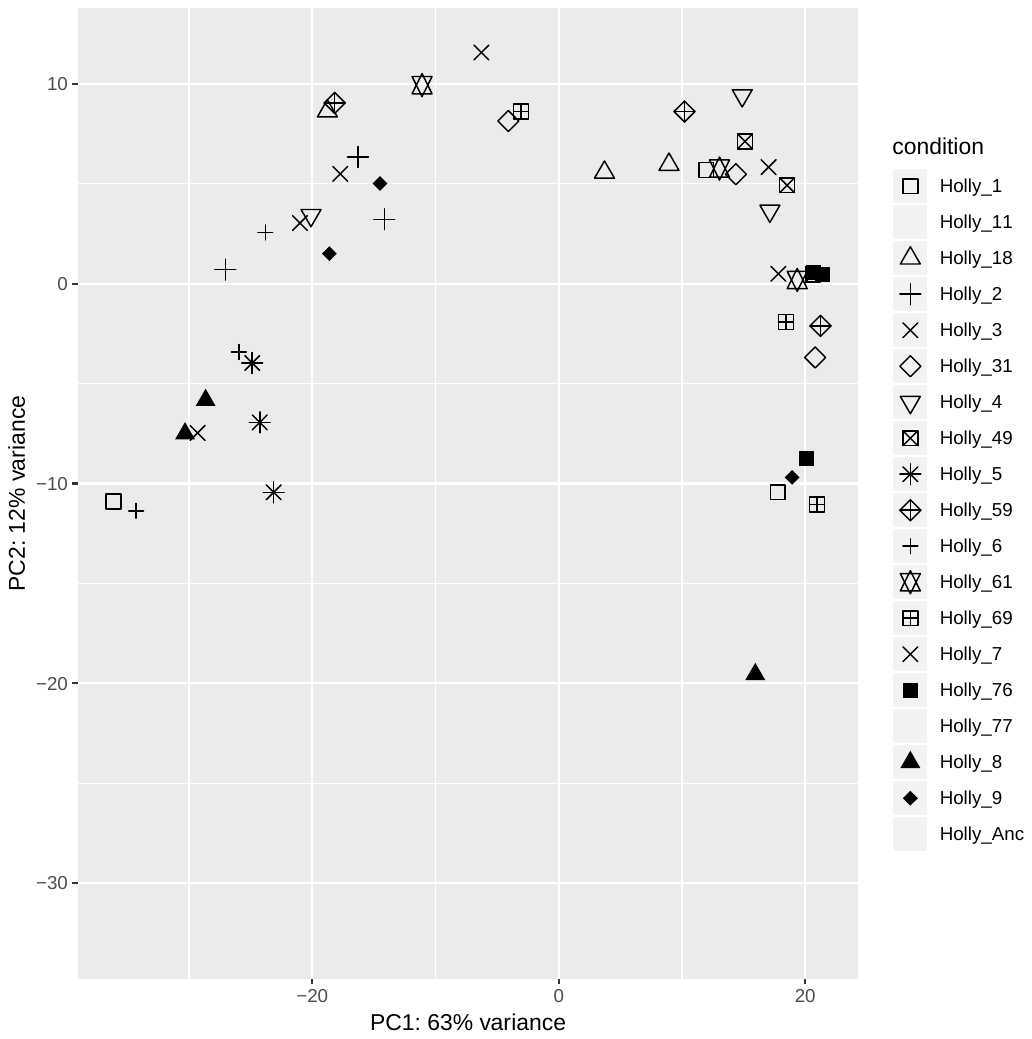

C: PCA of heterozygous ancestor sequencing run.

**Supplemental Figure 2:** There is no relationship between chromosome size and the number of aneuploidy (nondisjunction) events captured during mutation accumulation.

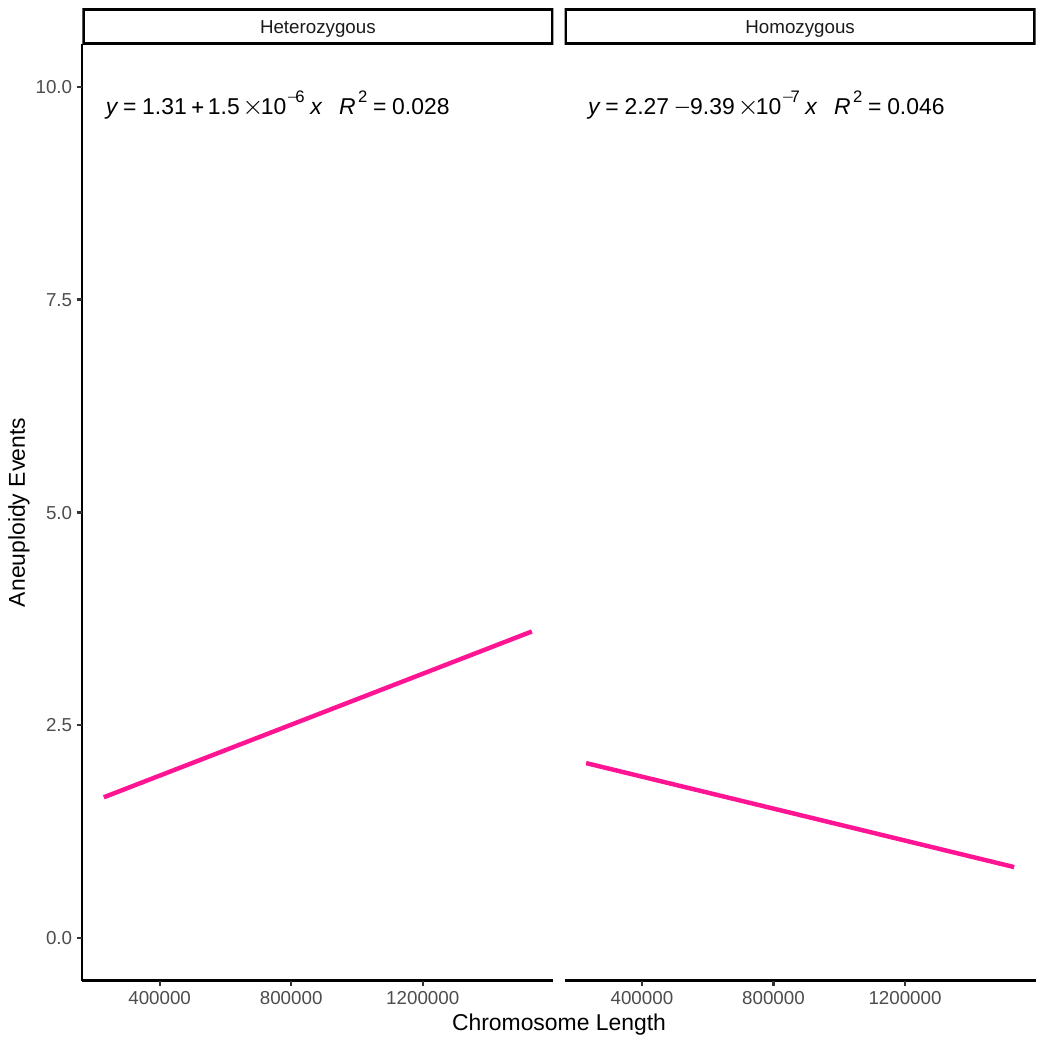

**Supplemental Figure 3:** Screenshot of Geneious alignment for SSD1 in S288C and our ancestral strains depicting a fully translated SSD1 protein.

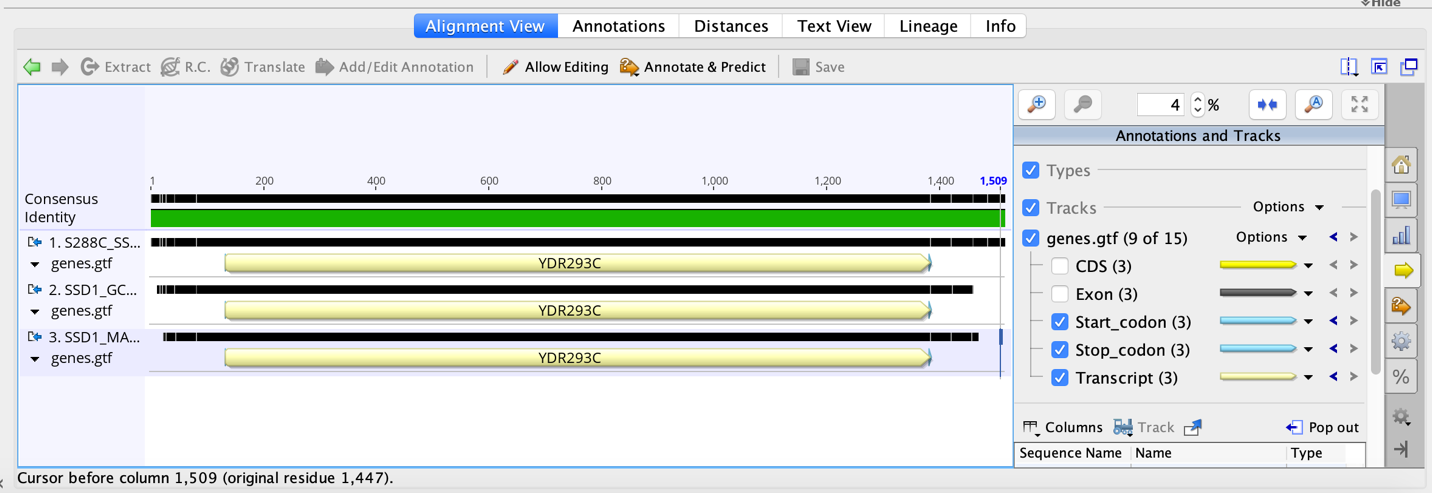

**Supplemental Figure 4:** Boxplots showing gene expression levels measured by log2(FPKMratio) for 16 heterozygous ancestor (first 16 columns) and 22 homozygous ancestor derived MA lines. Horizontal gray line is expectation if there is no change in gene expression. Dashed red line is expectation for 1.5-fold increase and dashed blue line is for 2-fold decrease in gene expression. Boxes in blue indicates that MA line is monosomic for the chromosome, red indicates trisomy, dark red indicates tetrasomy, pink indicates a partially duplicated chromosome, and gray indicates disomy (the normal state).

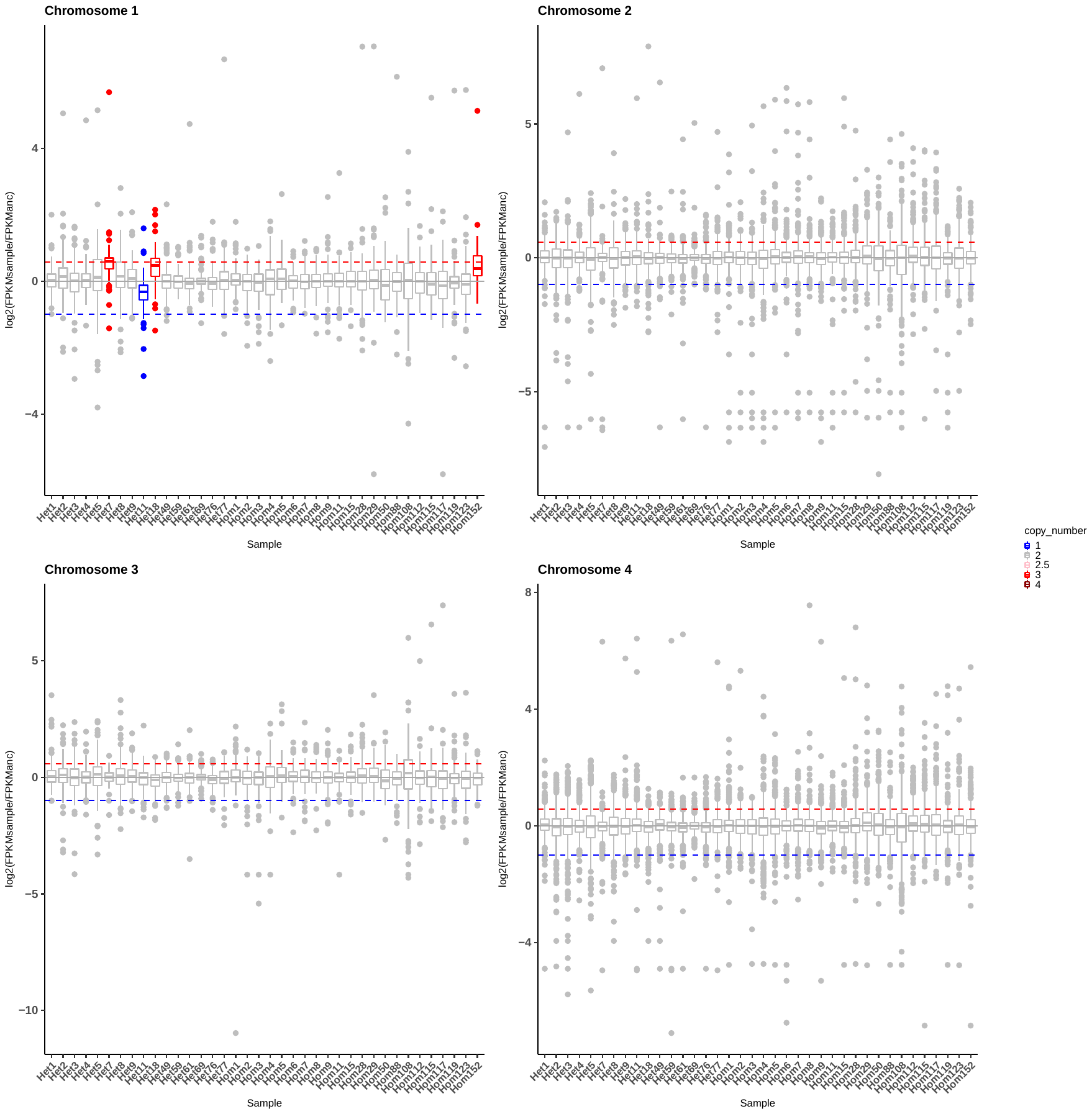

**Supplemental Figure 4 (cont)**

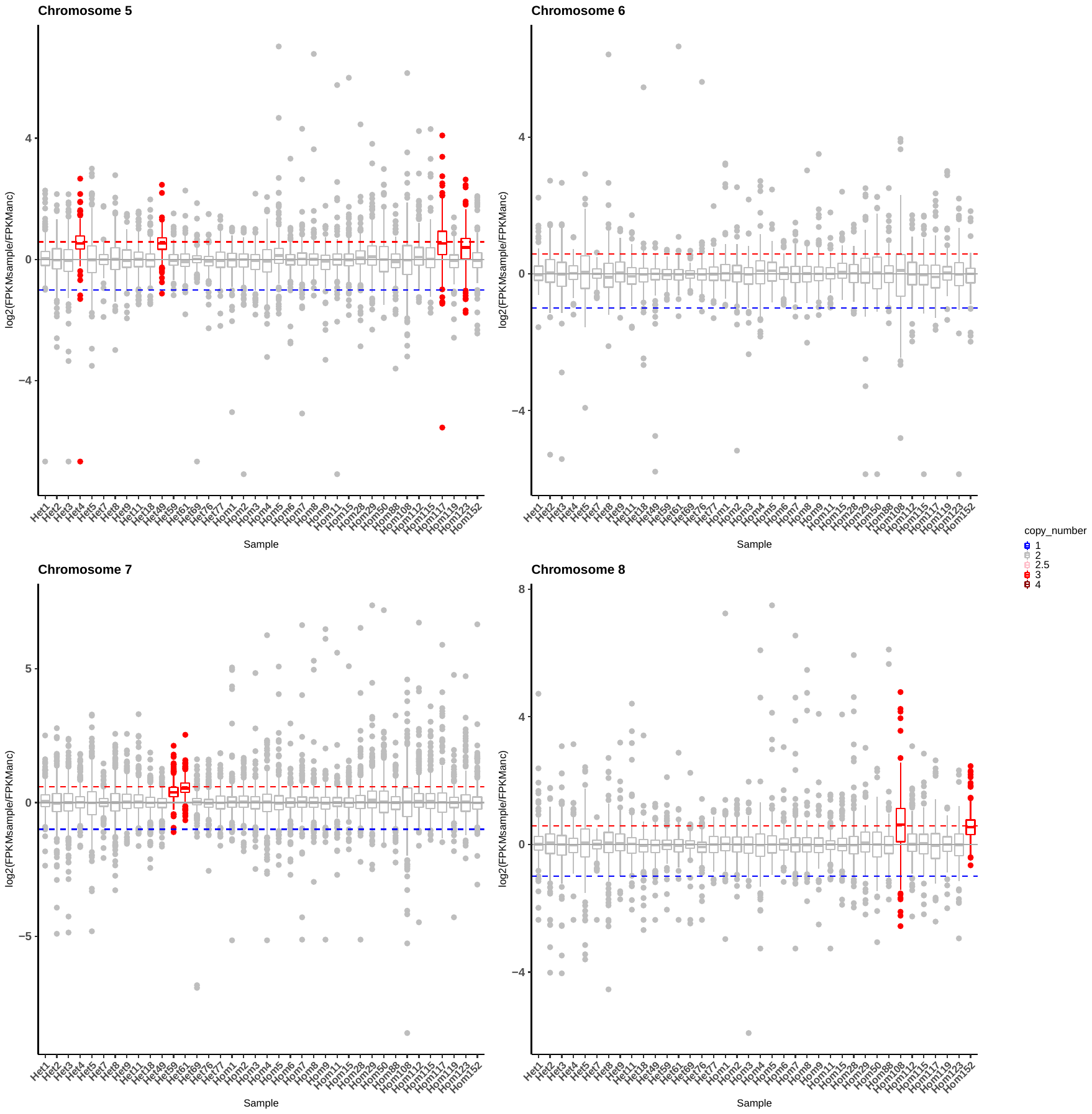

**Supplemental Figure 4 (cont)**

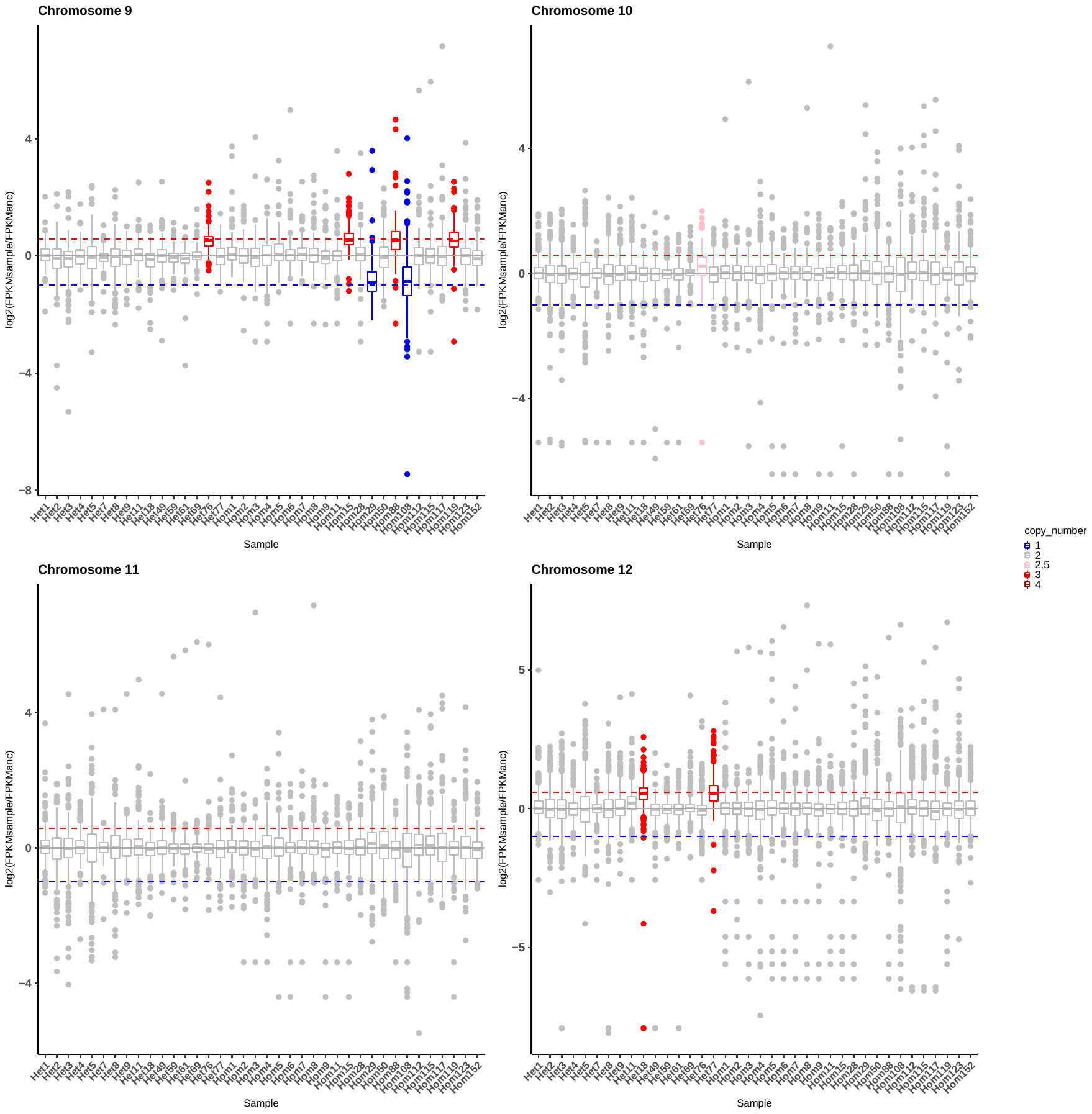

**Supplemental Figure 4 (cont)**

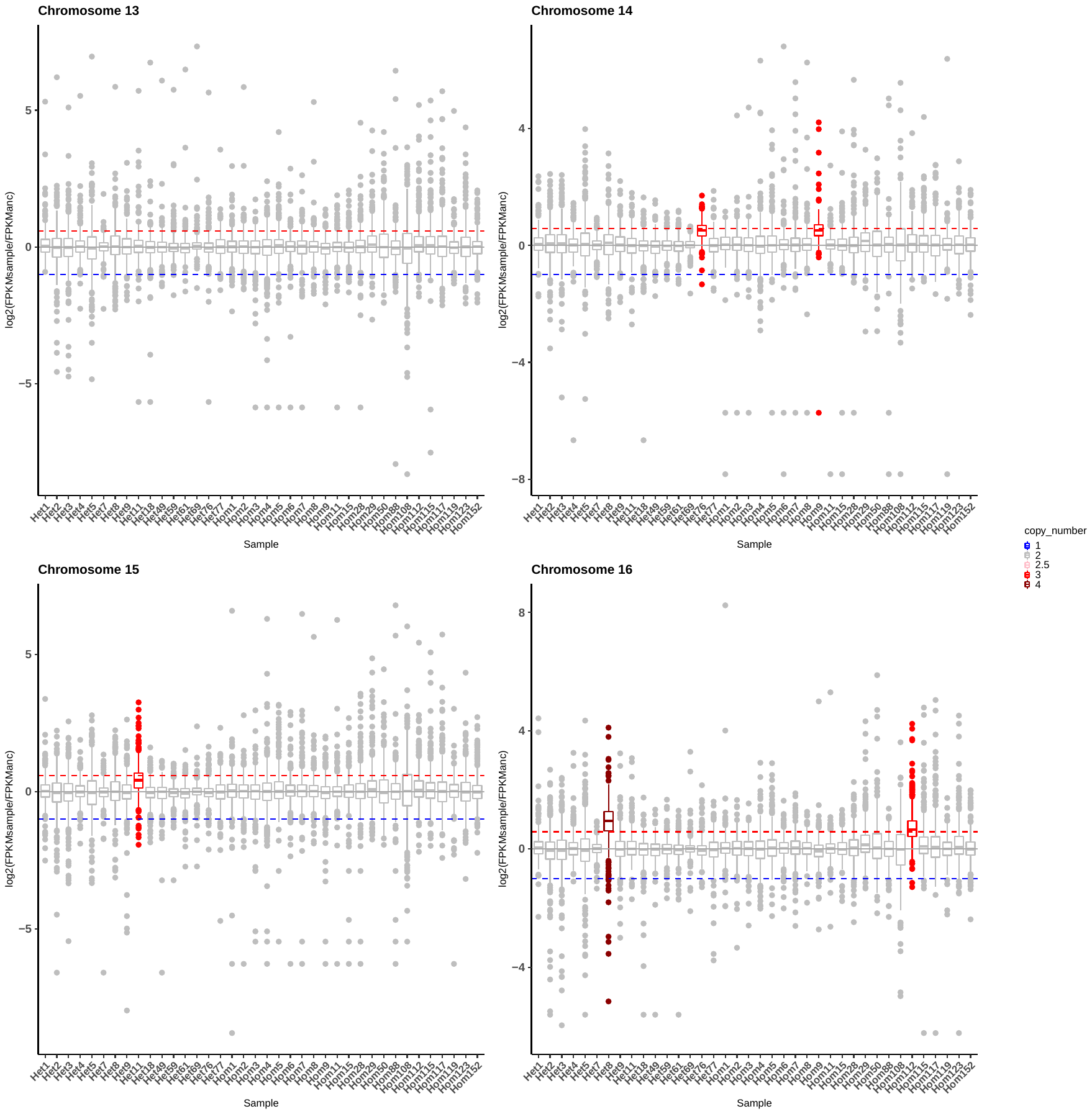

**Supplemental Figure 5:** Normalized count ratio distributions for genes in five euploid, heterozygous ancestor MA lines. Axes were cutoff at 4 even if there were more genes that had ratios greater than 4. Red dotted line: average ratio, blue dotted line: median ratio, black line: ratio of 1 (equal expression compared to ancestor). A; heterozygous line 1, B: heterozygous line 2, C: heterozygous line 3, D: heterozygous line 5, E: heterozygous line 9.

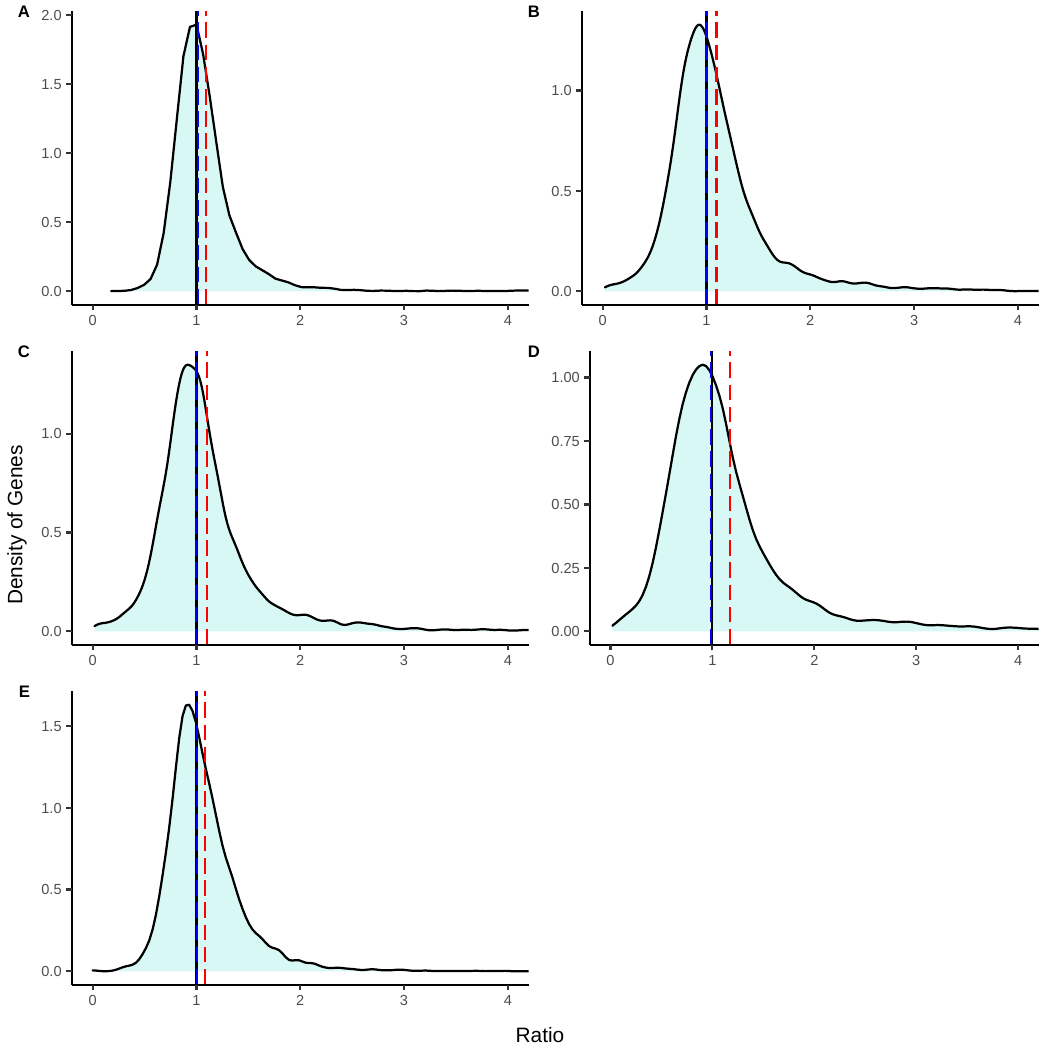

**Supplemental Figure 6:** Normalized count ratio distributions for genes in 9 euploid, homozygous ancestor MA lines. Genes with normalized count ratios that were 6 or greater are binned together. Red dotted line: average ratio, blue dotted line: median ratio, black line: ratio of 1 (equal expression compared to ancestor). A: homozygous line 1, B: homozygous line 2, C: homozygous line 3, D: homozygous line 4, E: homozygous line 5, F: homozygous line 7, G: homozygous line 8, H: homozygous line 50, I: homozygous line 115.

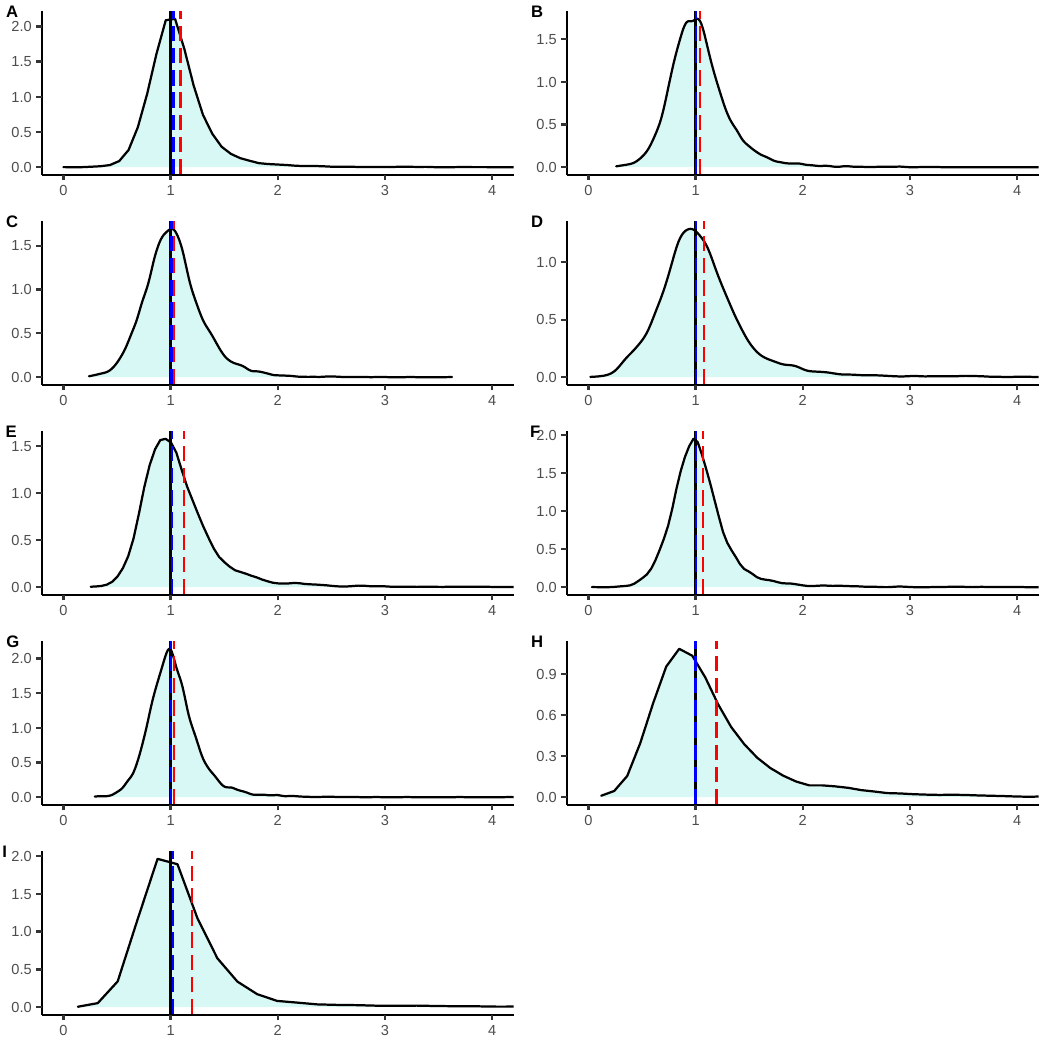

**Supplemental Figure 7:** Ratio distributions for aneuploid lines from the heterozygous ancestor. A-line 7 chromosome I, B-line 4 chromosome V, C- line 49 chr V, D-line 59 chr VII, E-line 61 chr VII, F-line 77 chr XII. Black line is expected ratio for euploid sample; blue line is the actual mean ratio of trans genes ; magenta line is the expected ratio for a trisomic chromosome; red line is the actual mean ratio for the given trisomic chromosome.

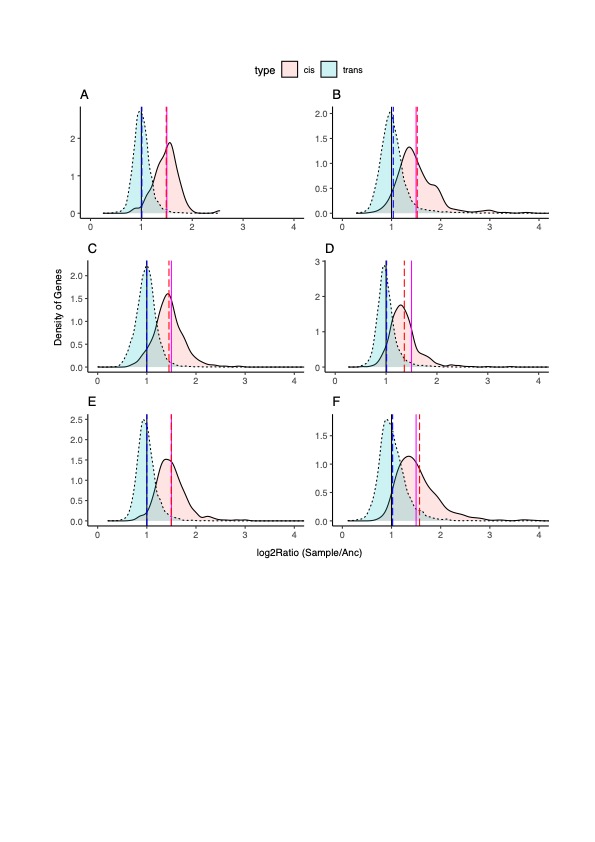

**Supplemental Figure 8:** Ratio distributions for trisomic lines from the homozygous ancestor. A: line 15 chr IX, B: line 88 chr IX, C: line 119 chr IX, D: line 9 chr XIV, E: line 112 chr XVI, F: line 117 chr V, G: line 123 chr V. Black line is expected ratio for euploid sample; cyan line is the actual mean ratio; magenta line is the expected ratio for a trisomic chromosome; red line is the actual mean ratio for the given trisomic chromosome.

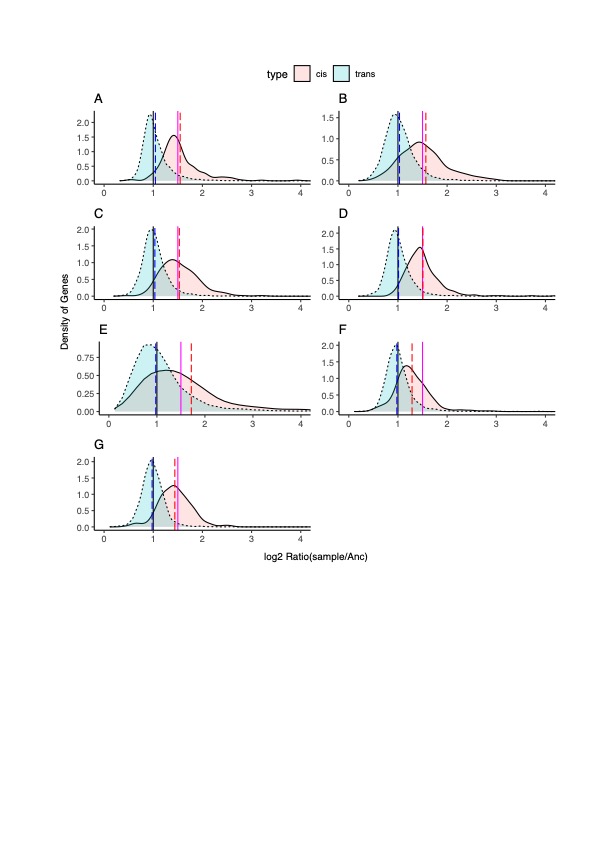

**Supplemental Figure 9:** Ratio distributions for lines from the heterozygous ancestor that have multiple aneuploidies. A: line 11 which is monosomic for chromosome I and trisomic for chromosome XV (red dashed line is the mean ratio for chromosome I and purple dashed line is the mean ratio for chromosome XV), B: line 18 which is trisomic for chromosome I and trisomic for chromosome XII (red dashed line is the mean ratio for chromosome I and purple dashed line is mean ratio for chromosome XII). For all samples: black line is expected ratio for euploid sample; yellow line is expected ratio of the trans genes in a line with the given aneuploid chromosome; cyan line is the actual mean ratio of the trans genes; magenta line is the expected ratio for a trisomic chromosome; green line is the expected ratio for a monosomic chromosome.

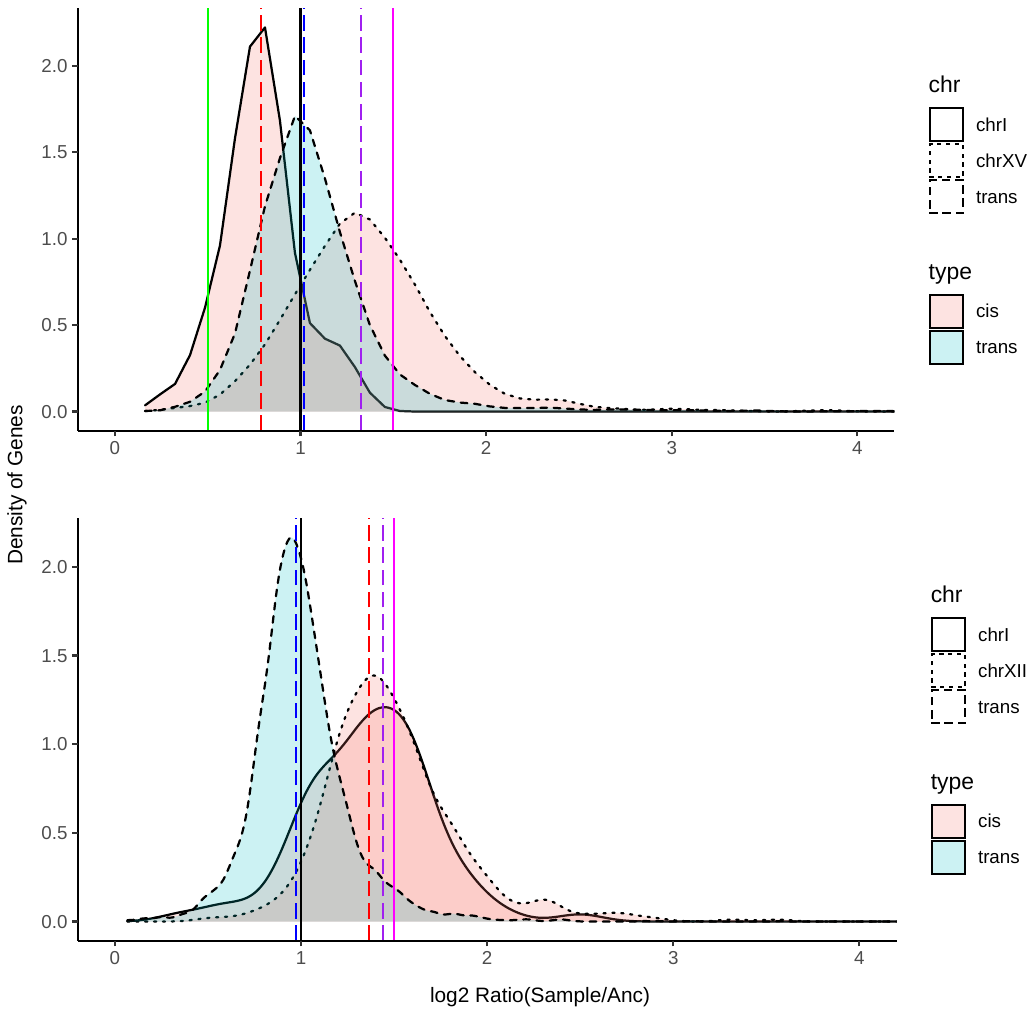

**Supplemental Figure 10:** Ratio distribution for homozygous ancestor samples with multiple aneuploidies. For all samples: black line is expected ratio for euploid sample; blue line is the actual mean ratio of the trans genes; magenta line is the expected ratio for a trisomic chromosome; green line is the expected ratio for a monosomic chromosome. A: Line 152, B: line 108.

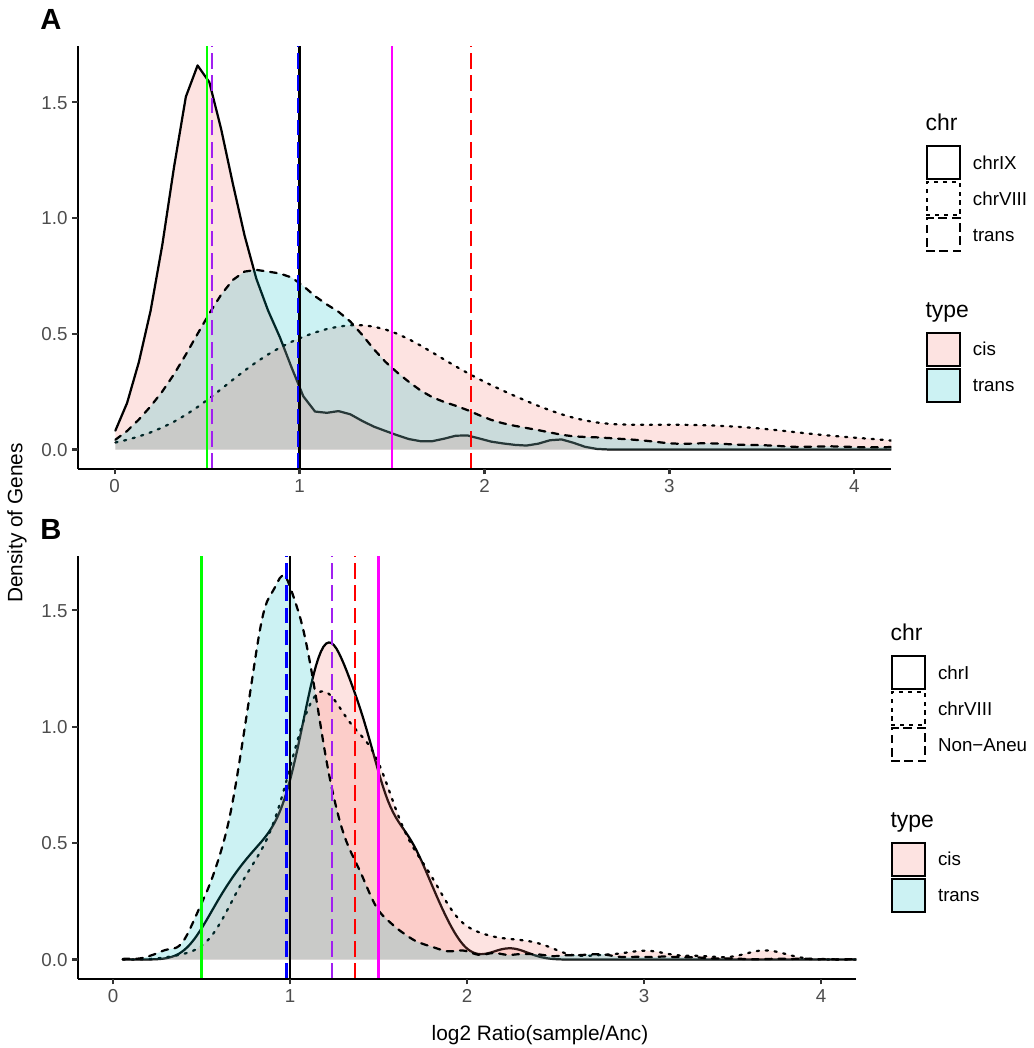

**Supplemental Figure 11:** Ratio distribution for heterozygous ancestor line 76, which has two trisomies and a partial duplication of chromosome X. Black line: expected ratio for a euploid line; blue line: mean ratio of trans genes; magenta line: expected for trisomy; red line: actual mean of trisomy IX, purple line: actual mean of trisomy XIV, pink line: actual mean of partial duplication chromosome X.

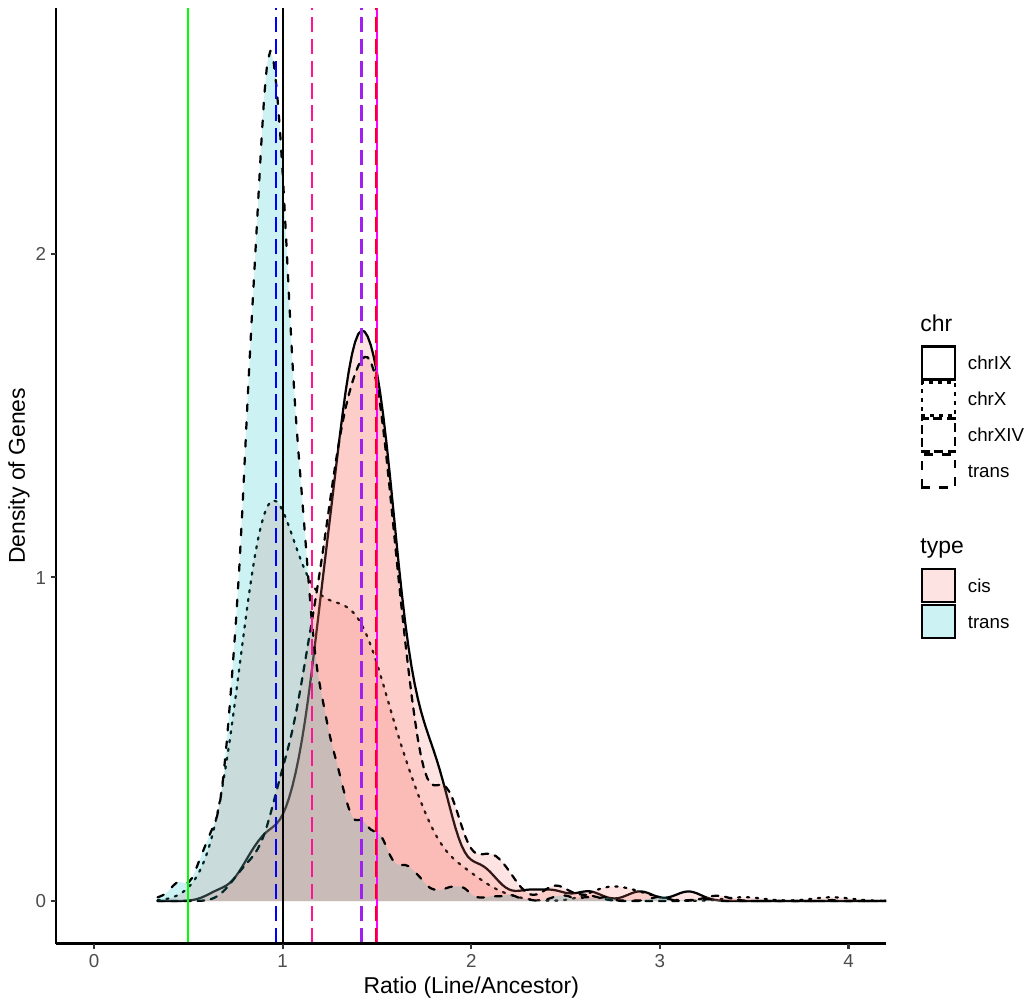

**Supplemental Figure 12:** Ratio distribution for homozygous ancestor line 29, which is monosomic for chromosome IX. Black line: expected ratio for a euploid line; blue line: mean ratio of trans genes; green line: expected for monosomy; red line: actual mean of chromosome IX.
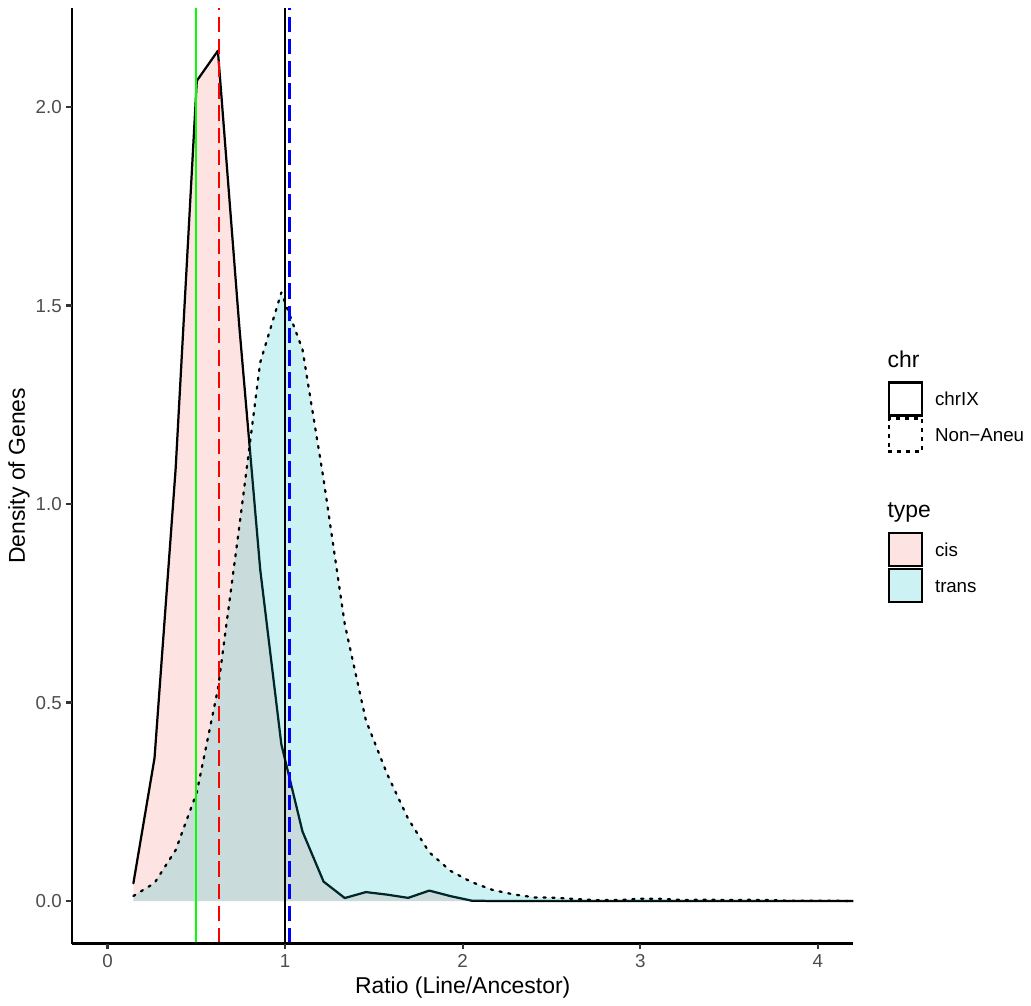

**Supplemental Figure 13:** Gene ontology analysis of commonly differentially expressed genes in the ESR from heterozygous ancestor euploid samples.
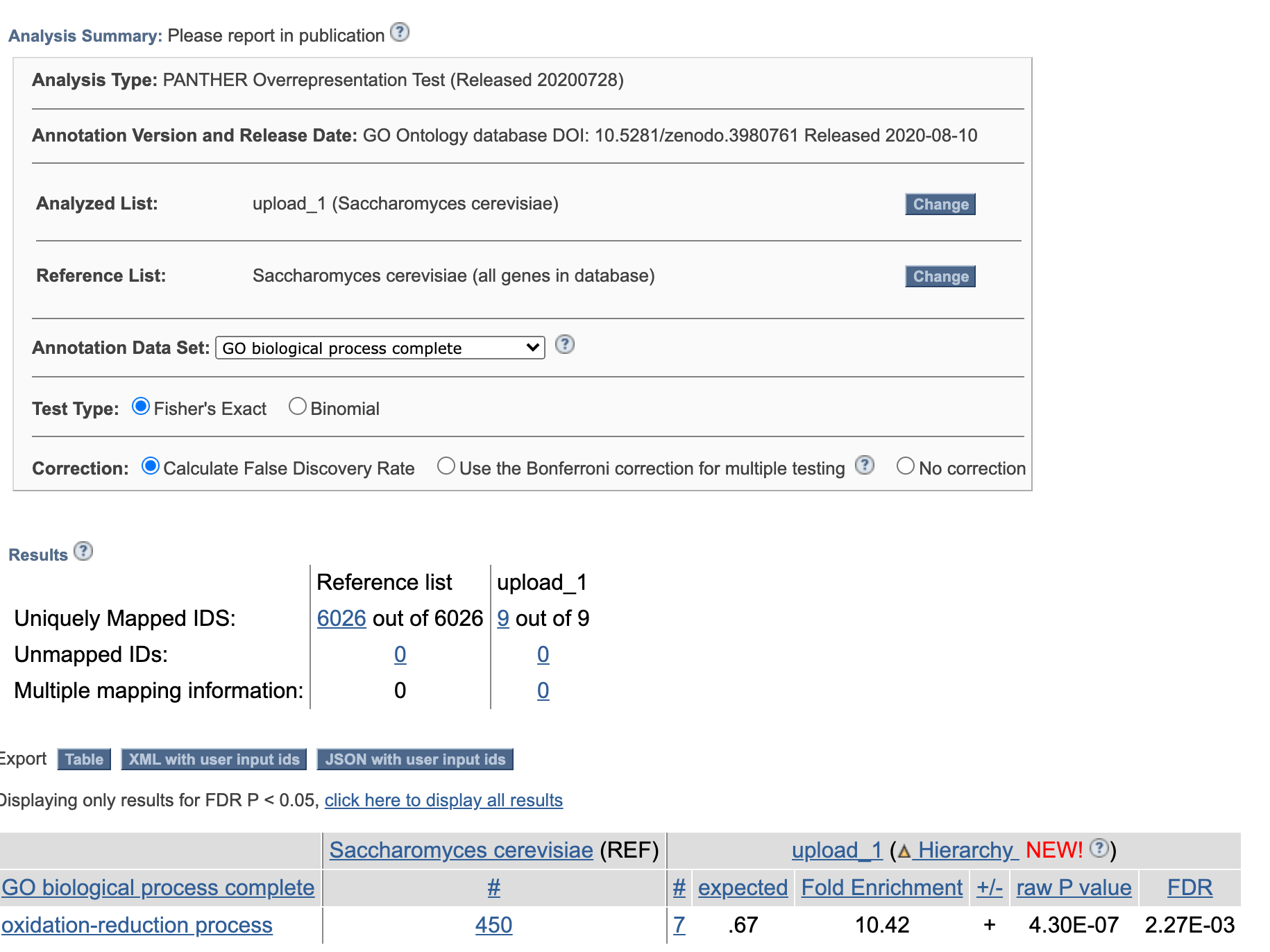

**Supplemental Table 1**. **Expected mean ratio of gene expression in MA lines trisomic for single chromosome shown (ratio of Line/Anc)**. Because more reads map to trisomic chromosome, number mapping to rest of genome will decline slightly.

| Trisomic Chromosome | Expected ratio for trans genes |
| --- | --- |
| 1 | 0.98 |
| 2 | 0.94 |
| 3 | 0.97 |
| 4 | 0.89 |
| 5 | 0.95 |
| 6 | 0.98 |
| 7 | 0.92 |
| 8 | 0.96 |
| 9 | 0.96 |
| 10 | 0.94 |
| 11 | 0.95 |
| 12 | 0.92 |
| 13 | 0.93 |
| 14 | 0.94 |
| 15 | 0.92 |
| 16 | 0.93 |

**Supplemental Table 2:** **Variance comparison between aneuploid and euploid chromosomes in aneuploid versus euploid MA lines.** Note: MA line Het-76 carries a partial duplication of chromosome 14. Since it is a partial duplication, we did not examine the variance in expression for that chromosome.

| MA line | | Aneuploidy present* | Euploid line comparison | | Aneuploid chromosomes  Levene's Statistic p-value | | Euploid chromosomes  Levene's Statistic p-value |
| --- | --- | --- | --- | --- | --- | --- | --- |
| Hom-152 | 1, 7 | | Hom-1 | Chr 1: 0.21  Chr 7: 0.58 | | 0.49 | |
| Hom-117 | 5 | | Hom-2 | 3.9E-11 | | 1.2E-12 | |
| Hom-123 | 5 | | Hom-3 | 3.5E-15 | | 6.5E-10 | |
| Hom-108 | 8, 9^m^ | | Hom-4 | Chr 8: 1.40E-09  Chr 9: 4.2E-3 | | 2.08E-56 | |
| Hom-15 | 9 | | Hom-5 | 0.15 | | 2.5E-4 | |
| Hom-29 | 9^m^ | | Hom-6 | 0.82 | | 2.1E-9 | |
| Hom-88 | 9 | | Hom-7 | 4.4E-4 | | 4.6E-10 | |
| Hom-119 | 9 | | Hom-8 | 2.1E-1 | | 0.19 | |
| Hom-9 | 14 | | Hom-11 | 0.0041 | | 7.8E-5 | |
| Hom-112 | 16 | | Hom-28 | 1.6E-5 | | 0.98 | |
| Het-7 | 1 | | Het-1 | 0.16 | | 6.8E-17 | |
| Het-11 | 1^m^, 15 | | Het-2 | Chr 1: 0.16  Chr 15: 0.11 | | 2.2E-8 | |
| Het-18 | 1, 12 | | Het-3 | Chr 1: 1.7E-4  Chr 12: 2.21E-10 | | 2.038E-02 | |
| Het-4 | 5 | | Het-5 | 2.3E-4 | | 1.5E-9 | |
| Het-49 | 5 | | Het-9 | 1.4E-6 | | 0.078 | |
| Het-59 | 7 | | Het-69 | 3.1E-30 | | 3.8E-9 | |
| Het-61 | 7 | | Het-1 | 0.023 | | 0.0024 | |
| Het-76 | 9, 10^p^, 14 | | Het-2 | Chr 9: 0.899  Chr 14: 0.0088 | | 4.55E-12 | |
| Het-77 | 12 | | Het-3 | 5.6E-13 | | 0.0032 | |
| Het-8 | 16^tet^ | | Het-5 | 0.12 | | 1.1E-16 | |

***** ^m^monosomic. ^p^partial duplication, ^tet^tetrasomic. All others trisomic.

**Supplemental Table 3: Expected gene expression for categories 1-6.** 1. Not dosage compensated: these genes have expression levels not significantly different as those predicted by their gene dose. 2. Partially dosage compensated: these genes show less-extreme gene expression changes than predicted by their dose. 3. Fully dosage compensated: these genes show no change in expression in response to changes in gene dose. 4. Over-dosage compensated: these genes show changes in expression that are in the opposite direction of the change in gene dose. 5. Anti-dosage compensated genes show more extreme changes in expression than predicted by the change in gene dose (in the direction of the aneuploidy – i.e. monosomic genes would have lower gene expression than predicted by monosomy).

The expected expression of a gene for each expression category following trisomy (2🡪3 copies) or monosomy (2🡪1copy).

|  | Expression Category | | | | |
| --- | --- | --- | --- | --- | --- |
| Copy number | 1 | 2 | 3 | 4 | 5 |
| 2 🡪 3 | 1.5x | 1-1.5x | 1x | <1x | > 1.5x |
| 2 🡪 1 | 0.5x | 0.5-1x | 1x | >1x | <0.5x |

**Supplemental Table 4:** Histone genes do not show evidence for dosage compensation.

| **Sample 9 HH: 3n Chrom XIV** | **CHROM** | **p val** |
| --- | --- | --- |
| YNL030W | chrXIV | 0.47345714 |
| YNL031C | chrXIV | 0.02953073 |
| **Sample 5 Hh: Euploid** |  |  |
| YNL031C | chrXIV | -0.5735449 |
| **Sample 6 Hh: Euploid** |  |  |
| YNL031C | chrXIV | -0.602327796 |
| **Sample 2 Hh: Euploid** |  |  |
| YNL031C | chrXIV | -0.783581586 |
| **Sample 4 HH: Euploid** |  |  |
| YNL031C | chrXIV | -0.571639056 |
| **Sample 76 Hh: 3n Chrom XIV** |  |  |
| YNL030W | chrXIV | 1.30707478 |
| **Sample 5 Hh: Euploid** |  |  |
| YNL030W | chrXIV | -0.456516426 |
| **Sample 6 Hh: Euploid** |  |  |
| YNL030W | chrXIV | -1.069145026 |
| **Sample 2 Hh: Euploid** |  |  |
| YNL030W | chrXIV | -1.163354563 |
| **Sample 5 HH: Euploid** |  |  |
| YNL030W | chrXIV | 1.260712151 |
| **Sample 8 HH: Euploid** |  |  |
| YNL030W | chrXIV | 1.015915942 |
| **Sample 76 Hh: 3n Chrom XIV** |  |  |
| YNL031C | chrXIV | 1.286739747 |
| **Sample 11 Hh:: 3n Chrom XV** |  |  |
| YOL012C | chrXV | 0.02705394 |
| **Sample 5 Hh: Euploid** |  |  |
| YOL012C | chrXV | -0.746267705 |
| **Sample 6 Hh:: Euploid** |  |  |
| YOL012C | chrXV | -0.661943487 |
| **Sample 2 Hh: Euploid** |  |  |
| YOL012C | chrXV | -0.867495188 |
| **Sample 112 HH: 3n Chrom XVI** |  |  |
| YPL127C | chrXVI | 0.60187799 |
| **Sample 5 Hh: Euploid** |  |  |
| YPL127C | chrXVI | -1.338035551 |
| **Sample 2 Hh: Euploid** |  |  |
| YPL127C | chrXVI | -1.031325188 |
| **Sample 7 HH: Euploid** |  |  |
| YPL127C | chrXVI | -0.599943193 |
| **Sample 8 Hh: 4n Chrom XVI** |  |  |
| YPL127C | chrXVI | -0.5278193 |
