## Supplementary material for "No evidence for whole-chromosome dosage compensation or global transcriptomic expression differences in spontaneously-aneuploid mutation accumulation lines of *Saccharomyces cerevisiae*": ANOVAs

**Supplemental Data:** ANOVA results for heterozygous ancestor lines

Call:

lm(formula = y ~ Line, data = chr10DataGC)

Residuals:

Min	1Q Median	3Q	Max

-5.8764 -0.2104 -0.0039 0.2181 2.6883

Coefficients:

Estimate Std. Error t value Pr(>|t|)

(Intercept) 0.043450	0.027753	1.566 0.117491

Line2	-0.071882	0.039248 -1.831 0.067080 .

Line3	-0.105918	0.039248 -2.699 0.006981 **

Line4	-0.043226	0.039248 -1.101 0.270788

Line5	-0.079235	0.039248 -2.019 0.043552 *

Line7	-0.055174	0.039248 -1.406 0.159845

Line8	-0.076579	0.039248 -1.951 0.051087 .

Line9	-0.013909	0.039248 -0.354 0.723060

Line11	-0.020249	0.039248 -0.516 0.605936

Line18	-0.130685	0.039248 -3.330 0.000875 ***

Line49	-0.092968	0.039248 -2.369 0.017882 *

Line59	-0.066252	0.039248 -1.688 0.091459 .

Line61	-0.087181	0.039248 -2.221 0.026370 *

Line69	-0.008323	0.039248 -0.212 0.832061

Line76	0.189169	0.039248	4.820 1.47e-06 ***

Line77	-0.051198	0.039248 -1.304 0.192124

---

Signif. codes: 0 '***' 0.001 '**' 0.01 '*' 0.05 '.' 0.1 ' ' 1

Residual standard error: 0.536 on 5952 degrees of freedom

Multiple R-squared: 0.01681,	Adjusted R-squared: 0.01434

F-statistic: 6.786 on 15 and 5952 DF, p-value: 8.272e-15

Call:

lm(formula = y ~ Line, data = chr11DataGC)

Residuals:

Min	1Q Median	3Q	Max

-4.0085 -0.2092 -0.0084 0.1985 6.0719

Coefficients:

Estimate Std. Error t value Pr(>|t|)

(Intercept) 0.07613	0.02930	2.599 0.00938 **

Line2	-0.12361	0.04143 -2.984 0.00286 **

Line3	-0.10789	0.04143 -2.604 0.00924 **

Line4	-0.04171	0.04143 -1.007 0.31417

Line5	-0.07830	0.04143 -1.890 0.05883 .

Line7	-0.06313	0.04143 -1.524 0.12764

Line8	-0.08273	0.04143 -1.997 0.04591 *

Line9	-0.07338	0.04143 -1.771 0.07660 .

Line11	-0.03711	0.04143 -0.896 0.37043

Line18	-0.13973	0.04143 -3.373 0.00075 ***

Line49	-0.06644	0.04143 -1.604 0.10886

Line59	-0.07867	0.04143 -1.899 0.05766 .

Line61	-0.09706	0.04143 -2.343 0.01919 *

Line69	-0.05870	0.04143 -1.417 0.15662

Line76	-0.07057	0.04143 -1.703 0.08859 .

Line77	-0.09405	0.04143 -2.270 0.02325 *

---

Signif. codes: 0 '***' 0.001 '**' 0.01 '*' 0.05 '.' 0.1 ' ' 1

Residual standard error: 0.5378 on 5376 degrees of freedom

Multiple R-squared: 0.00372,	Adjusted R-squared: 0.0009402

F-statistic: 1.338 on 15 and 5376 DF, p-value: 0.1696

Call:

lm(formula = y ~ Line, data = chr12DataGC)

Residuals:

Min	1Q Median	3Q	Max

-8.4229 -0.2265 -0.0140 0.2157 4.8866

Coefficients:

Estimate Std. Error t value Pr(>|t|)

(Intercept) 0.10028	0.02566	3.908 9.36e-05 ***

Line2	-0.05953	0.03629 -1.640 0.100945

Line3	-0.12219	0.03629 -3.367 0.000762 ***

Line4	-0.10250	0.03629 -2.825 0.004742 **

Line5	-0.07449	0.03629 -2.053 0.040127 *

Line7	-0.09646	0.03629 -2.658 0.007865 **

Line8	-0.09899	0.03629 -2.728 0.006386 **

Line9	-0.01809	0.03629 -0.498 0.618163

Line11	0.11652	0.03629	3.211 0.001327 **

Line18	0.41606	0.03629 11.466 < 2e-16 ***

Line49	-0.20237	0.03629 -5.577 2.52e-08 ***

Line59	-0.11217	0.03629 -3.091 0.002000 **

Line61	-0.15248	0.03629 -4.202 2.67e-05 ***

Line69	-0.05896	0.03629 -1.625 0.104219

Line76	-0.12915	0.03629 -3.559 0.000374 ***

Line77	0.49135	0.03629 13.541 < 2e-16 ***

---

Signif. codes: 0 '***' 0.001 '**' 0.01 '*' 0.05 '.' 0.1 ' ' 1

Residual standard error: 0.5884 on 8400 degrees of freedom

Multiple R-squared: 0.0944,	Adjusted R-squared: 0.09278

F-statistic: 58.37 on 15 and 8400 DF, p-value: < 2.2e-16

Call:

lm(formula = y ~ Line, data = chr13DataGC)

Residuals:

Min	1Q Median	3Q	Max

-5.7087 -0.2288 -0.0230 0.1967 7.2752

Coefficients:

Estimate Std. Error t value Pr(>|t|)

(Intercept) 0.10826	0.02623	4.127 3.71e-05 ***

Line2	-0.13847	0.03709 -3.733 0.000191 ***

Line3	-0.14548	0.03709 -3.922 8.87e-05 ***

Line4	-0.07265	0.03709 -1.959 0.050206 .

Line5	-0.12587	0.03709 -3.393 0.000694 ***

Line7	-0.10293	0.03709 -2.775 0.005538 **

Line8	-0.07761	0.03709 -2.092 0.036446 *

Line9	-0.07164	0.03709 -1.931 0.053483 .

Line11	-0.07257	0.03709 -1.956 0.050444 .

Line18	-0.13354	0.03709 -3.600 0.000320 ***

Line49	-0.11282	0.03709 -3.041 0.002363 **

Line59	-0.09163	0.03709 -2.470 0.013526 *

Line61	-0.12742	0.03709 -3.435 0.000595 ***

Line69	-0.05003	0.03709 -1.349 0.177508

Line76	-0.11462	0.03709 -3.090 0.002008 **

Line77	-0.09452	0.03709 -2.548 0.010849 *

---

Signif. codes: 0 '***' 0.001 '**' 0.01 '*' 0.05 '.' 0.1 ' ' 1

Residual standard error: 0.5753 on 7680 degrees of freedom

Multiple R-squared: 0.00407,	Adjusted R-squared: 0.002125

F-statistic: 2.092 on 15 and 7680 DF, p-value: 0.007888

Call:

lm(formula = y ~ Line, data = chr14DataGC)

Residuals:

Min	1Q Median	3Q	Max

-6.6706 -0.2183 0.0001 0.2052 3.8830

Coefficients:

Estimate Std. Error t value Pr(>|t|)

(Intercept) 0.08537	0.02413	3.538 0.000406 ***

Line2	-0.01535	0.03412 -0.450 0.652930

Line3	-0.01504	0.03412 -0.441 0.659520

Line4	-0.07725	0.03412 -2.264 0.023622 *

Line5	0.01185	0.03412	0.347 0.728506

Line7	-0.08029	0.03412 -2.353 0.018662 *

Line8	-0.03715	0.03412 -1.089 0.276401

Line9	-0.02938	0.03412 -0.861 0.389281

Line11	-0.10141	0.03412 -2.972 0.002973 **

Line18	-0.12089	0.03412 -3.543 0.000399 ***

Line49	-0.14967	0.03412 -4.386 1.17e-05 ***

Line59	-0.08370	0.03412 -2.453 0.014204 *

Line61	-0.10711	0.03412 -3.139 0.001704 **

Line69	-0.06939	0.03412 -2.034 0.042036 *

Line76	0.42708	0.03412 12.515 < 2e-16 ***

Line77	-0.07181	0.03412 -2.104 0.035380 *

---

Signif. codes: 0 '***' 0.001 '**' 0.01 '*' 0.05 '.' 0.1 ' ' 1

Residual standard error: 0.4921 on 6640 degrees of freedom

Multiple R-squared: 0.06223,	Adjusted R-squared: 0.06011

F-statistic: 29.37 on 15 and 6640 DF, p-value: < 2.2e-16

Call:

lm(formula = y ~ Line, data = chr15DataGC)

Residuals:

Min	1Q Median	3Q	Max

-7.9311 -0.2113 0.0074 0.2269 3.3413

Coefficients:

Estimate Std. Error t value Pr(>|t|)

(Intercept) 0.03715	0.02200	1.689 0.091262 .

Line2	-0.09965	0.03111 -3.204 0.001362 **

Line3	-0.10116	0.03111 -3.252 0.001149 **

Line4	-0.03825	0.03111 -1.230 0.218831

Line5	-0.05921	0.03111 -1.903 0.057013 .

Line7	-0.07368	0.03111 -2.369 0.017873 *

Line8	-0.03093	0.03111 -0.994 0.320127

Line9	-0.07430	0.03111 -2.388 0.016939 *

Line11	0.38144	0.03111 12.263 < 2e-16 ***

Line18	-0.08076	0.03111 -2.596 0.009442 **

Line49	-0.10819	0.03111 -3.478 0.000507 ***

Line59	-0.08489	0.03111 -2.729 0.006363 **

Line61	-0.10178	0.03111 -3.272 0.001071 **

Line69	-0.03643	0.03111 -1.171 0.241557

Line76	-0.07287	0.03111 -2.343 0.019172 *

Line77	-0.05099	0.03111 -1.639 0.101213

---

Signif. codes: 0 '***' 0.001 '**' 0.01 '*' 0.05 '.' 0.1 ' ' 1

Residual standard error: 0.5242 on 9072 degrees of freedom

Multiple R-squared: 0.04413,	Adjusted R-squared: 0.04255

F-statistic: 27.92 on 15 and 9072 DF, p-value: < 2.2e-16

Call:

lm(formula = y ~ Line, data = chr16DataGC)

Residuals:

Min	1Q Median	3Q	Max

-6.0310 -0.2075 0.0006 0.2120 4.3841

Coefficients:

Estimate Std. Error t value Pr(>|t|)

(Intercept) 0.07071	0.02519	2.807 0.005010 **

Line2	-0.14621	0.03562 -4.105 4.09e-05 ***

Line3	-0.12369	0.03562 -3.472 0.000519 ***

Line4	-0.02710	0.03562 -0.761 0.446880

Line5	-0.11288	0.03562 -3.169 0.001537 **

Line7	-0.06461	0.03562 -1.814 0.069741 .

Line8	0.80446	0.03562 22.584 < 2e-16 ***

Line9	-0.07048	0.03562 -1.979 0.047907 *

Line11	-0.03090	0.03562 -0.867 0.385771

Line18	-0.12181	0.03562 -3.419 0.000631 ***

Line49	-0.10457	0.03562 -2.936 0.003338 **

Line59	-0.07560	0.03562 -2.122 0.033851 *

Line61	-0.10737	0.03562 -3.014 0.002584 **

Line69	-0.04652	0.03562 -1.306 0.191580

Line76	-0.09823	0.03562 -2.758 0.005836 **

Line77	-0.09818	0.03562 -2.756 0.005861 **

---

Signif. codes: 0 '***' 0.001 '**' 0.01 '*' 0.05 '.' 0.1 ' ' 1

Residual standard error: 0.5518 on 7664 degrees of freedom

Multiple R-squared: 0.1353,	Adjusted R-squared: 0.1336

F-statistic: 79.93 on 15 and 7664 DF, p-value: < 2.2e-16

Call:

lm(formula = y ~ Line, data = chr1DataGC)

Residuals:

Min	1Q Median	3Q	Max

-3.9123 -0.2504 -0.0161 0.1929 6.5904

Coefficients:

Estimate Std. Error t value Pr(>|t|)

(Intercept) 0.048111	0.060519	0.795	0.427

Line2	0.104434	0.085587	1.220	0.223

Line3	-0.067564	0.085587 -0.789	0.430

Line4	0.023977	0.085587	0.280	0.779

Line5	0.062101	0.085587	0.726	0.468

Line7	0.498164	0.085587	5.821 7.05e-09 ***

Line8	0.088339	0.085587	1.032	0.302

Line9	0.055628	0.085587	0.650	0.516

Line11	-0.405439	0.085587 -4.737 2.36e-06 ***

Line18	0.384978	0.085587	4.498 7.34e-06 ***

Line49	-0.023784	0.085587 -0.278	0.781

Line59	-0.032118	0.085587 -0.375	0.708

Line61	-0.045047	0.085587 -0.526	0.599

Line69	-0.002338	0.085587 -0.027	0.978

Line76	-0.066835	0.085587 -0.781	0.435

Line77	0.043517	0.085587	0.508	0.611

---

Signif. codes: 0 '***' 0.001 '**' 0.01 '*' 0.05 '.' 0.1 ' ' 1

Residual standard error: 0.6142 on 1632 degrees of freedom

Multiple R-squared: 0.08813,	Adjusted R-squared: 0.07975

F-statistic: 10.51 on 15 and 1632 DF, p-value: < 2.2e-16

Call:

lm(formula = y ~ Line, data = chr2DataGC)

Residuals:

Min	1Q Median	3Q	Max

-7.0775 -0.2115 0.0029 0.2187 7.9057

Coefficients:

Estimate Std. Error t value Pr(>|t|)

(Intercept) 0.017668	0.028117	0.628	0.530

Line2	-0.048387	0.039764 -1.217	0.224

Line3	-0.059357	0.039764 -1.493	0.136

Line4	-0.011688	0.039764 -0.294	0.769

Line5	-0.050971	0.039764 -1.282	0.200

Line7	-0.032734	0.039764 -0.823	0.410

Line8	-0.033988	0.039764 -0.855	0.393

Line9	-0.014371	0.039764 -0.361	0.718

Line11	0.015613	0.039764	0.393	0.695

Line18	-0.033646	0.039764 -0.846	0.398

Line49	-0.042802	0.039764 -1.076	0.282

Line59	-0.039665	0.039764 -0.998	0.319

Line61	-0.052958	0.039764 -1.332	0.183

Line69	0.018538	0.039764	0.466	0.641

Line76	-0.064105	0.039764 -1.612	0.107

Line77	-0.007319	0.039764 -0.184	0.854

Residual standard error: 0.5824 on 6848 degrees of freedom

Multiple R-squared: 0.001848,	Adjusted R-squared: -0.0003383

F-statistic: 0.8453 on 15 and 6848 DF, p-value: 0.6271

Call:

lm(formula = y ~ Line, data = chr3DataGC)

Residuals:

Min	1Q Median	3Q	Max

-4.1443 -0.2437 -0.0002 0.2351 3.4290

Coefficients:

Estimate Std. Error t value Pr(>|t|)

(Intercept) 0.094173	0.039757	2.369 0.017919 *

Line2	-0.028822	0.056225 -0.513 0.608258

Line3	-0.106175	0.056225 -1.888 0.059080 .

Line4	-0.061657	0.056225 -1.097 0.272908

Line5	-0.007464	0.056225 -0.133 0.894392

Line7	-0.096421	0.056225 -1.715 0.086476 .

Line8	0.001606	0.056225	0.029 0.977209

Line9	-0.013958	0.056225 -0.248 0.803956

Line11	-0.156085	0.056225 -2.776 0.005540 **

Line18	-0.236045	0.056225 -4.198 2.78e-05 ***

Line49	-0.128924	0.056225 -2.293 0.021924 *

Line59	-0.111442	0.056225 -1.982 0.047571 *

Line61	-0.136831	0.056225 -2.434 0.015012 *

Line69	-0.082590	0.056225 -1.469 0.141968

Line76	-0.201466	0.056225 -3.583 0.000345 ***

Line77	-0.109070	0.056225 -1.940 0.052497 .

---

Signif. codes: 0 '***' 0.001 '**' 0.01 '*' 0.05 '.' 0.1 ' ' 1

Residual standard error: 0.5168 on 2688 degrees of freedom

Multiple R-squared: 0.0177,	Adjusted R-squared: 0.01222

F-statistic: 3.229 on 15 and 2688 DF, p-value: 2.428e-05

Call:

lm(formula = y ~ Line, data = chr4DataGC)

Residuals:

Min	1Q Median	3Q	Max

-7.0639 -0.2002 -0.0024 0.1963 6.6083

Coefficients:

Estimate Std. Error t value Pr(>|t|)

(Intercept) 0.05219	0.01735	3.008 0.002632 **

Line2	-0.09727	0.02453 -3.965 7.38e-05 ***

Line3	-0.08061	0.02453 -3.286 0.001020 **

Line4	-0.06206	0.02453 -2.530 0.011429 *

Line5	-0.06643	0.02453 -2.708 0.006780 **

Line7	-0.07875	0.02453 -3.210 0.001331 **

Line8	-0.06269	0.02453 -2.556 0.010615 *

Line9	-0.04707	0.02453 -1.919 0.055058 .

Line11	-0.01139	0.02453 -0.464 0.642351

Line18	-0.09985	0.02453 -4.070 4.73e-05 ***

Line49	-0.04351	0.02453 -1.774 0.076141 .

Line59	-0.08559	0.02453 -3.489 0.000487 ***

Line61	-0.09692	0.02453 -3.951 7.84e-05 ***

Line69	-0.04154	0.02453 -1.693 0.090469 .

Line76	-0.08729	0.02453 -3.558 0.000375 ***

Line77	-0.06242	0.02453 -2.544 0.010957 *

---

Signif. codes: 0 '***' 0.001 '**' 0.01 '*' 0.05 '.' 0.1 ' ' 1

Residual standard error: 0.4904 on 12768 degrees of freedom

Multiple R-squared: 0.003389,	Adjusted R-squared: 0.002218

F-statistic: 2.895 on 15 and 12768 DF, p-value: 0.000138

Call:

lm(formula = y ~ Line, data = chr5DataGC)

Residuals:

Min	1Q Median	3Q	Max

-7.2076 -0.2305 -0.0114 0.2243 2.9353

Coefficients:

Estimate Std. Error t value Pr(>|t|)

(Intercept) 0.07353	0.03048	2.412 0.01589 *

Line2	-0.05512	0.04311 -1.279 0.20111

Line3	-0.09706	0.04311 -2.251 0.02441 *

Line4	0.47164	0.04311 10.940 < 2e-16 ***

Line5	-0.01202	0.04311 -0.279 0.78042

Line7	-0.08066	0.04311 -1.871 0.06142 .

Line8	-0.04762	0.04311 -1.105 0.26935

Line9	-0.04835	0.04311 -1.122 0.26210

Line11	-0.06920	0.04311 -1.605 0.10851

Line18	-0.09618	0.04311 -2.231 0.02573 *

Line49	0.43073	0.04311	9.991 < 2e-16 ***

Line59	-0.09199	0.04311 -2.134 0.03290 *

Line61	-0.11620	0.04311 -2.695 0.00705 **

Line69	-0.08027	0.04311 -1.862 0.06267 .

Line76	-0.14149	0.04311 -3.282 0.00104 **

Line77	-0.08140	0.04311 -1.888 0.05908 .

---

Signif. codes: 0 '***' 0.001 '**' 0.01 '*' 0.05 '.' 0.1 ' ' 1

Residual standard error: 0.5306 on 4832 degrees of freedom

Multiple R-squared: 0.1002,	Adjusted R-squared: 0.09739

F-statistic: 35.87 on 15 and 4832 DF, p-value: < 2.2e-16

Call:

lm(formula = y ~ Line, data = chr6DataGC)

Residuals:

Min	1Q Median	3Q	Max

-5.6564 -0.2300 -0.0128 0.2017 6.6094

Coefficients:

Estimate Std. Error t value Pr(>|t|)

(Intercept) 0.060758	0.053282	1.140	0.2543

Line2	0.010294	0.075352	0.137	0.8913

Line3	-0.073950	0.075352 -0.981	0.3265

Line4	-0.047476	0.075352 -0.630	0.5287

Line5	0.006771	0.075352	0.090	0.9284

Line7	-0.066170	0.075352 -0.878	0.3800

Line8	-0.023501	0.075352 -0.312	0.7552

Line9	0.008857	0.075352	0.118	0.9064

Line11	-0.131298	0.075352 -1.742	0.0816 .

Line18	-0.079830	0.075352 -1.059	0.2895

Line49	-0.189594	0.075352 -2.516	0.0119 *

Line59	-0.084888	0.075352 -1.127	0.2601

Line61	-0.019293	0.075352 -0.256	0.7979

Line69	-0.064961	0.075352 -0.862	0.3887

Line76	-0.028100	0.075352 -0.373	0.7093

Line77	-0.048529	0.075352 -0.644	0.5196

---

Signif. codes: 0 '***' 0.001 '**' 0.01 '*' 0.05 '.' 0.1 ' ' 1

Residual standard error: 0.5909 on 1952 degrees of freedom

Multiple R-squared: 0.007905,	Adjusted R-squared: 0.0002812

F-statistic: 1.037 on 15 and 1952 DF, p-value: 0.413

Call:

lm(formula = y ~ Line, data = chr7DataGC)

Residuals:

Min	1Q Median	3Q	Max

-6.9356 -0.2159 -0.0103 0.2103 3.2824

Coefficients:

Estimate Std. Error t value Pr(>|t|)

(Intercept) 0.08642	0.02124	4.069 4.76e-05 ***

Line2	-0.08930	0.03003 -2.974 0.002952 **

Line3	-0.09023	0.03003 -3.005 0.002667 **

Line4	-0.05858	0.03003 -1.951 0.051123 .

Line5	-0.07834	0.03003 -2.609 0.009109 **

Line7	-0.08903	0.03003 -2.964 0.003041 **

Line8	-0.07072	0.03003 -2.355 0.018559 *

Line9	-0.03257	0.03003 -1.084 0.278217

Line11	-0.01526	0.03003 -0.508 0.611400

Line18	-0.09476	0.03003 -3.155 0.001609 **

Line49	-0.11943	0.03003 -3.977 7.04e-05 ***

Line59	0.32743	0.03003 10.902 < 2e-16 ***

Line61	0.47031	0.03003 15.660 < 2e-16 ***

Line69	-0.07586	0.03003 -2.526 0.011558 *

Line76	-0.10798	0.03003 -3.595 0.000326 ***

Line77	-0.07257	0.03003 -2.416 0.015698 *

---

Signif. codes: 0 '***' 0.001 '**' 0.01 '*' 0.05 '.' 0.1 ' ' 1

Residual standard error: 0.5021 on 8928 degrees of freedom

Multiple R-squared: 0.09279,	Adjusted R-squared: 0.09126

F-statistic: 60.88 on 15 and 8928 DF, p-value: < 2.2e-16

Call:

lm(formula = y ~ Line, data = chr8DataGC)

Residuals:

Min	1Q Median	3Q	Max

-4.5644 -0.2176 0.0048 0.2149 4.6639

Coefficients:

Estimate Std. Error t value Pr(>|t|)

(Intercept) 0.05886	0.03105	1.896 0.05802 .

Line2	-0.06366	0.04391 -1.450 0.14713

Line3	-0.07981	0.04391 -1.818 0.06917 .

Line4	-0.07171	0.04391 -1.633 0.10250

Line5	-0.01711	0.04391 -0.390 0.69686

Line7	-0.07331	0.04391 -1.670 0.09505 .

Line8	-0.03550	0.04391 -0.809 0.41877

Line9	0.01237	0.04391	0.282 0.77810

Line11	-0.03954	0.04391 -0.901 0.36783

Line18	-0.11449	0.04391 -2.608 0.00914 **

Line49	-0.13559	0.04391 -3.088 0.00203 **

Line59	-0.06524	0.04391 -1.486 0.13739

Line61	-0.08516	0.04391 -1.940 0.05249 .

Line69	-0.05358	0.04391 -1.220 0.22237

Line76	-0.14273	0.04391 -3.251 0.00116 **

Line77	-0.04720	0.04391 -1.075 0.28238

---

Signif. codes: 0 '***' 0.001 '**' 0.01 '*' 0.05 '.' 0.1 ' ' 1

Residual standard error: 0.5241 on 4544 degrees of freedom

Multiple R-squared: 0.006497,	Adjusted R-squared: 0.003217

F-statistic: 1.981 on 15 and 4544 DF, p-value: 0.01321

Call:

lm(formula = y ~ Line, data = chr9DataGC)

Residuals:

Min	1Q Median	3Q	Max

-5.1851 -0.2101 0.0108 0.2193 2.5443

Coefficients:

Estimate Std. Error t value Pr(>|t|)

(Intercept) 0.0270074 0.0304687	0.886 0.375461

Line2	-0.1472850 0.0430892 -3.418 0.000637 ***

Line3	-0.1627409 0.0430892 -3.777 0.000161 ***

Line4	-0.0589257 0.0430892 -1.368 0.171544

Line5	-0.0939892 0.0430892 -2.181 0.029227 *

Line7	-0.0942619 0.0430892 -2.188 0.028762 *

Line8	-0.0987465 0.0430892 -2.292 0.021981 *

Line9	-0.0661724 0.0430892 -1.536 0.124697

Line11	-0.0005015 0.0430892 -0.012 0.990715

Line18	-0.1986916 0.0430892 -4.611 4.14e-06 ***

Line49	-0.0353929 0.0430892 -0.821 0.411480

Line59	-0.0890176 0.0430892 -2.066 0.038909 *

Line61	-0.1136020 0.0430892 -2.636 0.008414 **

Line69	-0.0200296 0.0430892 -0.465 0.642074

Line76	0.5054586 0.0430892 11.731 < 2e-16 ***

Line77	-0.0612492 0.0430892 -1.421 0.155271

---

Signif. codes: 0 '***' 0.001 '**' 0.01 '*' 0.05 '.' 0.1 ' ' 1

Residual standard error: 0.4621 on 3664 degrees of freedom

Multiple R-squared: 0.09833,	Adjusted R-squared: 0.09464

F-statistic: 26.64 on 15 and 3664 DF, p-value: < 2.2e-16

Supplemental Data: ANOVA results for homozygous ancestor lines.

Call:

lm(formula = y ~ Line, data = chr1DataMA)

Residuals:

Min	1Q Median	3Q	Max

-5.3734 -0.3819 -0.0722 0.2750 6.9357

Coefficients:

Estimate Std. Error t value Pr(>|t|)

(Intercept) 0.13417	0.07986	1.680 0.093365 .

Line50	-0.13759	0.11294 -1.218 0.223524

Line112	-0.08817	0.11294 -0.781 0.435231

Line115	-0.10362	0.11294 -0.918 0.359163

Line117	-0.21546	0.11294 -1.908 0.056816 .

Line123	-0.12842	0.11294 -1.137 0.255866

Line152	0.40447	0.11294	3.581 0.000365 ***

---

Signif. codes: 0 '***' 0.001 '**' 0.01 '*' 0.05 '.' 0.1 ' ' 1

Residual standard error: 0.8105 on 714 degrees of freedom

Multiple R-squared: 0.05276,	Adjusted R-squared: 0.0448

F-statistic: 6.629 on 6 and 714 DF, p-value: 7.792e-07

Call:

lm(formula = y ~ Line, data = chr2DataMA)

Residuals:

Min	1Q Median	3Q	Max

-7.4584 -0.2692 0.0081 0.2810 5.9203

Coefficients:

Estimate Std. Error t value Pr(>|t|)

(Intercept) 0.0129161 0.0373682	0.346	0.7296

Line2	-0.0553960 0.0528466 -1.048	0.2946

Line3	-0.0841277 0.0528466 -1.592	0.1114

Line4	-0.1156188 0.0528466 -2.188	0.0287 *

Line5	0.0712320 0.0528466	1.348	0.1777

Line7	0.0197163 0.0528466	0.373	0.7091

Line8	0.0117606 0.0528466	0.223	0.8239

Line9	-0.0737775 0.0528466 -1.396	0.1627

Line11	-0.0238928 0.0528466 -0.452	0.6512

Line15	0.0278875 0.0528466	0.528	0.5977

Line28	0.0446541 0.0528466	0.845	0.3981

Line29	0.0384835 0.0528466	0.728	0.4665

Line50	-0.0470287 0.0528466 -0.890	0.3735

Line88	-0.0459765 0.0528466 -0.870	0.3843

Line108	-0.1213815 0.0528466 -2.297	0.0216 *

Line112	0.0936873 0.0528466	1.773	0.0763 .

Line115	0.0196062 0.0528466	0.371	0.7106

Line117	-0.0000458 0.0528466 -0.001	0.9993

Line119	-0.0743308 0.0528466 -1.407	0.1596

Line123	-0.0247336 0.0528466 -0.468	0.6398

Line152	-0.0512348 0.0528466 -0.970	0.3323

---

Signif. codes: 0 '***' 0.001 '**' 0.01 '*' 0.05 '.' 0.1 ' ' 1

Residual standard error: 0.774 on 8988 degrees of freedom

Multiple R-squared: 0.005476,	Adjusted R-squared: 0.003263

F-statistic: 2.474 on 20 and 8988 DF, p-value: 0.0002697

Call:

lm(formula = y ~ Line, data = chr3DataMA)

Residuals:

Min	1Q	Median	3Q	Max

-10.9942 -0.2947	0.0014	0.3006	7.8459

Coefficients:

Estimate Std. Error t value Pr(>|t|)

(Intercept) 0.019637	0.052724	0.372	0.7096

Line2	-0.079572	0.074563 -1.067	0.2860

Line3	-0.125993	0.074563 -1.690	0.0912 .

Line4	-0.013303	0.074563 -0.178	0.8584

Line5	0.096657	0.074563	1.296	0.1949

Line7	-0.007711	0.074563 -0.103	0.9176

Line8	-0.017923	0.074563 -0.240	0.8101

Line9	-0.043570	0.074563 -0.584	0.5590

Line11	-0.072688	0.074563 -0.975	0.3297

Line15	-0.003437	0.074563 -0.046	0.9632

Line28	0.083700	0.074563	1.123	0.2617

Line29	0.101905	0.074563	1.367	0.1718

Line50	-0.096868	0.074563 -1.299	0.1940

Line88	-0.095966	0.074563 -1.287	0.1982

Line108	0.065116	0.074563	0.873	0.3826

Line112	-0.042174	0.074563 -0.566	0.5717

Line115	-0.033274	0.074563 -0.446	0.6554

Line117	-0.089899	0.074563 -1.206	0.2280

Line119	-0.023888	0.074563 -0.320	0.7487

Line123	-0.117804	0.074563 -1.580	0.1142

Line152	-0.068622	0.074563 -0.920	0.3575

---

Signif. codes: 0 '***' 0.001 '**' 0.01 '*' 0.05 '.' 0.1 ' ' 1

Residual standard error: 0.6854 on 3528 degrees of freedom

Multiple R-squared: 0.009427,	Adjusted R-squared: 0.003812

F-statistic: 1.679 on 20 and 3528 DF, p-value: 0.0297

Call:

lm(formula = y ~ Line, data = chr4DataMA)

Residuals:

Min	1Q Median	3Q	Max

-7.4796 -0.2538 0.0013 0.2542 7.5309

Coefficients:

Estimate Std. Error t value Pr(>|t|)

(Intercept) 0.0236639 0.0201918	1.172 0.24123

Line2	-0.0266121 0.0285555 -0.932 0.35138

Line3	-0.0656131 0.0285555 -2.298 0.02159 *

Line4	-0.0587243 0.0285555 -2.056 0.03975 *

Line5	-0.0202708 0.0285555 -0.710 0.47779

Line7	-0.0098373 0.0285555 -0.344 0.73048

Line8	0.0054828 0.0285555	0.192 0.84774

Line9	-0.0913638 0.0285555 -3.200 0.00138 **

Line11	-0.0153717 0.0285555 -0.538 0.59037

Line15	-0.0515555 0.0285555 -1.805 0.07102 .

Line28	0.0040727 0.0285555	0.143 0.88659

Line29	0.0726426 0.0285555	2.544 0.01097 *

Line50	0.0233862 0.0285555	0.819 0.41281

Line88	-0.0721350 0.0285555 -2.526 0.01154 *

Line108	-0.0856368 0.0285555 -2.999 0.00271 **

Line112	0.0555600 0.0285555	1.946 0.05171 .

Line115	0.0275490 0.0285555	0.965 0.33468

Line117	-0.0002823 0.0285555 -0.010 0.99211

Line119	-0.0396279 0.0285555 -1.388 0.16523

Line123	-0.0209933 0.0285555 -0.735 0.46224

Line152	-0.0705933 0.0285555 -2.472 0.01344 *

---

Signif. codes: 0 '***' 0.001 '**' 0.01 '*' 0.05 '.' 0.1 ' ' 1

Residual standard error: 0.5708 on 16758 degrees of freedom

Multiple R-squared: 0.005856,	Adjusted R-squared: 0.004669

F-statistic: 4.935 on 20 and 16758 DF, p-value: 2.365e-12

Call:

lm(formula = y ~ Line, data = chr5DataMA)

Residuals:

Min	1Q Median	3Q	Max

-7.0796 -0.2753 0.0103 0.2698 6.8648

Coefficients:

Estimate Std. Error t value Pr(>|t|)

(Intercept) -0.053867	0.037331 -1.443	0.1491

Line2	-0.008935	0.052794 -0.169	0.8656

Line3	-0.033448	0.052794 -0.634	0.5264

Line4	-0.032250	0.052794 -0.611	0.5413

Line5	0.213992	0.052794	4.053 5.11e-05 ***

Line7	0.057902	0.052794	1.097	0.2728

Line8	0.086235	0.052794	1.633	0.1024

Line9	-0.040397	0.052794 -0.765	0.4442

Line11	0.053846	0.052794	1.020	0.3078

Line15	0.078736	0.052794	1.491	0.1359

Line28	0.100687	0.052794	1.907	0.0565 .

Line29	0.128717	0.052794	2.438	0.0148 *

Line50	0.037280	0.052794	0.706	0.4801

Line88	-0.014415	0.052794 -0.273	0.7848

Line108	0.014469	0.052794	0.274	0.7840

Line112	0.147911	0.052794	2.802	0.0051 **

Line115	0.075746	0.052794	1.435	0.1514

Line117	0.535707	0.052794 10.147 < 2e-16 ***

Line119	-0.020049	0.052794 -0.380	0.7041

Line123	0.357888	0.052794	6.779 1.32e-11 ***

Line152	-0.020876	0.052794 -0.395	0.6925

---

Signif. codes: 0 '***' 0.001 '**' 0.01 '*' 0.05 '.' 0.1 ' ' 1

Residual standard error: 0.6498 on 6342 degrees of freedom

Multiple R-squared: 0.04339,	Adjusted R-squared: 0.04038

F-statistic: 14.38 on 20 and 6342 DF, p-value: < 2.2e-16

Call:

lm(formula = y ~ Line, data = chr5DataMA)

Residuals:

Min	1Q Median	3Q	Max

-7.0796 -0.2753 0.0103 0.2698 6.8648

Coefficients:

Estimate Std. Error t value Pr(>|t|)

(Intercept) -0.053867	0.037331 -1.443	0.1491

Line2	-0.008935	0.052794 -0.169	0.8656

Line3	-0.033448	0.052794 -0.634	0.5264

Line4	-0.032250	0.052794 -0.611	0.5413

Line5	0.213992	0.052794	4.053 5.11e-05 ***

Line7	0.057902	0.052794	1.097	0.2728

Line8	0.086235	0.052794	1.633	0.1024

Line9	-0.040397	0.052794 -0.765	0.4442

Line11	0.053846	0.052794	1.020	0.3078

Line15	0.078736	0.052794	1.491	0.1359

Line28	0.100687	0.052794	1.907	0.0565 .

Line29	0.128717	0.052794	2.438	0.0148 *

Line50	0.037280	0.052794	0.706	0.4801

Line88	-0.014415	0.052794 -0.273	0.7848

Line108	0.014469	0.052794	0.274	0.7840

Line112	0.147911	0.052794	2.802	0.0051 **

Line115	0.075746	0.052794	1.435	0.1514

Line117	0.535707	0.052794 10.147 < 2e-16 ***

Line119	-0.020049	0.052794 -0.380	0.7041

Line123	0.357888	0.052794	6.779 1.32e-11 ***

Line152	-0.020876	0.052794 -0.395	0.6925

---

Signif. codes: 0 '***' 0.001 '**' 0.01 '*' 0.05 '.' 0.1 ' ' 1

Residual standard error: 0.6498 on 6342 degrees of freedom

Multiple R-squared: 0.04339,	Adjusted R-squared: 0.04038

F-statistic: 14.38 on 20 and 6342 DF, p-value: < 2.2e-16

Call:

lm(formula = y ~ Line, data = chr6DataMA)

Residuals:

Min	1Q Median	3Q	Max

-5.3621 -0.3027 -0.0049 0.2581 3.9222

Coefficients:

Estimate Std. Error t value Pr(>|t|)

(Intercept) 0.108431	0.062052	1.747	0.0807 .

Line2	-0.129085	0.087754 -1.471	0.1414

Line3	-0.153480	0.087754 -1.749	0.0804 .

Line4	-0.058092	0.087754 -0.662	0.5080

Line5	-0.014594	0.087754 -0.166	0.8679

Line7	-0.103712	0.087754 -1.182	0.2374

Line8	-0.106896	0.087754 -1.218	0.2233

Line9	-0.032061	0.087754 -0.365	0.7149

Line11	-0.086260	0.087754 -0.983	0.3257

Line15	0.002664	0.087754	0.030	0.9758

Line28	-0.093758	0.087754 -1.068	0.2854

Line29	-0.046956	0.087754 -0.535	0.5926

Line50	-0.032637	0.087754 -0.372	0.7100

Line88	-0.090652	0.087754 -1.033	0.3017

Line108	-0.081861	0.087754 -0.933	0.3510

Line112	-0.089905	0.087754 -1.025	0.3057

Line115	-0.120181	0.087754 -1.370	0.1710

Line117	-0.103711	0.087754 -1.182	0.2374

Line119	-0.007226	0.087754 -0.082	0.9344

Line123	-0.069580	0.087754 -0.793	0.4279

Line152	-0.141146	0.087754 -1.608	0.1079

---

Signif. codes: 0 '***' 0.001 '**' 0.01 '*' 0.05 '.' 0.1 ' ' 1

Residual standard error: 0.6882 on 2562 degrees of freedom

Multiple R-squared: 0.004419,	Adjusted R-squared: -0.003353

F-statistic: 0.5686 on 20 and 2562 DF, p-value: 0.9356

Call:

lm(formula = y ~ Line, data = chr7DataMA)

Residuals:

Min	1Q Median	3Q	Max

-8.6286 -0.2708 -0.0183 0.2388 7.2742

Coefficients:

Estimate Std. Error t value Pr(>|t|)

(Intercept) 0.06604	0.02742	2.408	0.0161 *

Line2	-0.02973	0.03878 -0.767	0.4433

Line3	-0.05234	0.03878 -1.350	0.1771

Line4	-0.07429	0.03878 -1.916	0.0554 .

Line5	-0.02810	0.03878 -0.725	0.4687

Line7	-0.02971	0.03878 -0.766	0.4436

Line8	-0.04427	0.03878 -1.141	0.2537

Line9	-0.07003	0.03878 -1.806	0.0710 .

Line11	-0.05168	0.03878 -1.333	0.1827

Line15	-0.04035	0.03878 -1.040	0.2982

Line28	-0.01507	0.03878 -0.389	0.6976

Line29	0.03339	0.03878	0.861	0.3893

Line50	-0.02525	0.03878 -0.651	0.5150

Line88	-0.07270	0.03878 -1.875	0.0609 .

Line108	-0.05322	0.03878 -1.372	0.1700

Line112	0.03127	0.03878	0.806	0.4200

Line115	-0.02382	0.03878 -0.614	0.5391

Line117	-0.04131	0.03878 -1.065	0.2868

Line119	-0.06196	0.03878 -1.598	0.1101

Line123	-0.06786	0.03878 -1.750	0.0802 .

Line152	-0.09111	0.03878 -2.349	0.0188 *

---

Signif. codes: 0 '***' 0.001 '**' 0.01 '*' 0.05 '.' 0.1 ' ' 1

Residual standard error: 0.6484 on 11718 degrees of freedom

Multiple R-squared: 0.002374,	Adjusted R-squared: 0.0006718

F-statistic: 1.395 on 20 and 11718 DF, p-value: 0.1123

Call:

lm(formula = y ~ Line, data = chr8DataMA)

Residuals:

Min	1Q Median	3Q	Max

-5.8911 -0.2930 -0.0259 0.2594 7.4064

Coefficients:

Estimate Std. Error t value Pr(>|t|)

(Intercept) 0.042601	0.038278	1.113	0.2658

Line2	-0.027706	0.054134 -0.512	0.6088

Line3	-0.060945	0.054134 -1.126	0.2603

Line4	-0.063369	0.054134 -1.171	0.2418

Line5	0.044807	0.054134	0.828	0.4079

Line7	-0.010838	0.054134 -0.200	0.8413

Line8	-0.003586	0.054134 -0.066	0.9472

Line9	-0.051800	0.054134 -0.957	0.3387

Line11	-0.046988	0.054134 -0.868	0.3854

Line15	-0.025058	0.054134 -0.463	0.6435

Line28	0.008869	0.054134	0.164	0.8699

Line29	-0.007671	0.054134 -0.142	0.8873

Line50	-0.089729	0.054134 -1.658	0.0975 .

Line88	-0.026332	0.054134 -0.486	0.6267

Line108	0.584627	0.054134 10.800	<2e-16 ***

Line112	0.004753	0.054134	0.088	0.9300

Line115	-0.048766	0.054134 -0.901	0.3677

Line117	-0.128824	0.054134 -2.380	0.0174 *

Line119	-0.053209	0.054134 -0.983	0.3257

Line123	-0.120320	0.054134 -2.223	0.0263 *

Line152	0.500959	0.054134	9.254	<2e-16 ***

---

Signif. codes: 0 '***' 0.001 '**' 0.01 '*' 0.05 '.' 0.1 ' ' 1

Residual standard error: 0.6462 on 5964 degrees of freedom

Multiple R-squared: 0.06896,	Adjusted R-squared: 0.06584

F-statistic: 22.09 on 20 and 5964 DF, p-value: < 2.2e-16

Call:

lm(formula = y ~ Line, data = chr9DataMA)

Residuals:

Min	1Q Median	3Q	Max

-6.6107 -0.2948 -0.0375 0.2452 7.5298

Coefficients:

Estimate Std. Error t value Pr(>|t|)

(Intercept) 0.14399	0.04189	3.438 0.000592 ***

Line2	-0.13341	0.05924 -2.252 0.024363 *

Line3	-0.17489	0.05924 -2.952 0.003169 **

Line4	-0.10577	0.05924 -1.786 0.074236 .

Line5	0.02021	0.05924	0.341 0.733042

Line7	-0.06519	0.05924 -1.100 0.271200

Line8	-0.07968	0.05924 -1.345 0.178635

Line9	-0.15104	0.05924 -2.550 0.010813 *

Line11	-0.12438	0.05924 -2.100 0.035807 *

Line15	0.45026	0.05924	7.601 3.52e-14 ***

Line28	-0.06830	0.05924 -1.153 0.248981

Line29	-1.01011	0.05924 -17.052 < 2e-16 ***

Line50	-0.17944	0.05924 -3.029 0.002466 **

Line88	0.40900	0.05924	6.904 5.70e-12 ***

Line108	-0.98399	0.05924 -16.611 < 2e-16 ***

Line112	-0.10504	0.05924 -1.773 0.076246 .

Line115	-0.13069	0.05924 -2.206 0.027415 *

Line117	-0.09210	0.05924 -1.555 0.120076

Line119	0.40545	0.05924	6.844 8.64e-12 ***

Line123	-0.13884	0.05924 -2.344 0.019128 *

Line152	-0.21804	0.05924 -3.681 0.000235 ***

---

Signif. codes: 0 '***' 0.001 '**' 0.01 '*' 0.05 '.' 0.1 ' ' 1

Residual standard error: 0.6353 on 4809 degrees of freedom

Multiple R-squared: 0.2272,	Adjusted R-squared: 0.224

F-statistic: 70.68 on 20 and 4809 DF, p-value: < 2.2e-16

Call:

lm(formula = y ~ Line, data = chr10DataMA)

Residuals:

Min	1Q Median	3Q	Max

-6.4665 -0.2639 -0.0052 0.2582 7.2217

Coefficients:

Estimate Std. Error t value Pr(>|t|)

(Intercept) 0.029150	0.034072	0.856	0.392

Line2	-0.019959	0.048185 -0.414	0.679

Line3	-0.036097	0.048185 -0.749	0.454

Line4	-0.074748	0.048185 -1.551	0.121

Line5	0.001790	0.048185	0.037	0.970

Line7	-0.034818	0.048185 -0.723	0.470

Line8	0.005026	0.048185	0.104	0.917

Line9	-0.074034	0.048185 -1.536	0.124

Line11	0.014936	0.048185	0.310	0.757

Line15	0.017669	0.048185	0.367	0.714

Line28	-0.013893	0.048185 -0.288	0.773

Line29	0.054126	0.048185	1.123	0.261

Line50	-0.002511	0.048185 -0.052	0.958

Line88	-0.078711	0.048185 -1.634	0.102

Line108	-0.116090	0.048185 -2.409	0.016 *

Line112	0.061408	0.048185	1.274	0.203

Line115	0.048483	0.048185	1.006	0.314

Line117	-0.017116	0.048185 -0.355	0.722

Line119	-0.064576	0.048185 -1.340	0.180

Line123	-0.041089	0.048185 -0.853	0.394

Line152	-0.067294	0.048185 -1.397	0.163

---

Signif. codes: 0 '***' 0.001 '**' 0.01 '*' 0.05 '.' 0.1 ' ' 1

Residual standard error: 0.658 on 7812 degrees of freedom

Multiple R-squared: 0.004941,	Adjusted R-squared: 0.002394

F-statistic: 1.94 on 20 and 7812 DF, p-value: 0.00718

Call:

lm(formula = y ~ Line, data = chr11DataMA)

Residuals:

Min	1Q Median	3Q	Max

-5.3800 -0.2654 -0.0053 0.2548 7.1361

Coefficients:

Estimate Std. Error t value Pr(>|t|)

(Intercept) 0.046100	0.034231	1.347 0.17811

Line2	-0.069521	0.048411 -1.436 0.15102

Line3	-0.079502	0.048411 -1.642 0.10058

Line4	-0.041052	0.048411 -0.848 0.39647

Line5	-0.002335	0.048411 -0.048 0.96154

Line7	-0.035455	0.048411 -0.732 0.46396

Line8	-0.011645	0.048411 -0.241 0.80992

Line9	-0.124375	0.048411 -2.569 0.01021 *

Line11	-0.045212	0.048411 -0.934 0.35038

Line15	-0.051030	0.048411 -1.054 0.29187

Line28	0.004407	0.048411	0.091 0.92747

Line29	0.081709	0.048411	1.688 0.09149 .

Line50	0.040056	0.048411	0.827 0.40803

Line88	-0.102530	0.048411 -2.118 0.03422 *

Line108	-0.137665	0.048411 -2.844 0.00447 **

Line112	0.047871	0.048411	0.989 0.32277

Line115	0.037727	0.048411	0.779 0.43583

Line117	0.018711	0.048411	0.387 0.69913

Line119	-0.071063	0.048411 -1.468 0.14217

Line123	-0.012930	0.048411 -0.267 0.78940

Line152	-0.105463	0.048411 -2.179 0.02940 *

---

Signif. codes: 0 '***' 0.001 '**' 0.01 '*' 0.05 '.' 0.1 ' ' 1

Residual standard error: 0.6284 on 7056 degrees of freedom

Multiple R-squared: 0.008612,	Adjusted R-squared: 0.005802

F-statistic: 3.065 on 20 and 7056 DF, p-value: 4.792e-06

Call:

lm(formula = y ~ Line, data = chr12DataMA)

Residuals:

Min	1Q Median	3Q	Max

-7.9937 -0.2588 0.0086 0.2757 7.2950

Coefficients:

Estimate Std. Error t value Pr(>|t|)

(Intercept) 0.006544	0.032968	0.198	0.8427

Line2	-0.016368	0.046624 -0.351	0.7255

Line3	-0.055019	0.046624 -1.180	0.2380

Line4	-0.099940	0.046624 -2.144	0.0321 *

Line5	0.040056	0.046624	0.859	0.3903

Line7	-0.030373	0.046624 -0.651	0.5148

Line8	0.018685	0.046624	0.401	0.6886

Line9	-0.030881	0.046624 -0.662	0.5078

Line11	-0.004223	0.046624 -0.091	0.9278

Line15	-0.015575	0.046624 -0.334	0.7384

Line28	-0.015843	0.046624 -0.340	0.7340

Line29	0.061630	0.046624	1.322	0.1862

Line50	-0.063730	0.046624 -1.367	0.1717

Line88	-0.042324	0.046624 -0.908	0.3640

Line108	-0.005635	0.046624 -0.121	0.9038

Line112	0.025041	0.046624	0.537	0.5912

Line115	-0.025703	0.046624 -0.551	0.5814

Line117	-0.099652	0.046624 -2.137	0.0326 *

Line119	-0.034883	0.046624 -0.748	0.4544

Line123	-0.057875	0.046624 -1.241	0.2145

Line152	-0.030041	0.046624 -0.644	0.5194

---

Signif. codes: 0 '***' 0.001 '**' 0.01 '*' 0.05 '.' 0.1 ' ' 1

Residual standard error: 0.7561 on 11025 degrees of freedom

Multiple R-squared: 0.002738,	Adjusted R-squared: 0.000929

F-statistic: 1.514 on 20 and 11025 DF, p-value: 0.06584

Call:

lm(formula = y ~ Line, data = chr13DataMA)

Residuals:

Min	1Q Median	3Q	Max

-8.2370 -0.2719 -0.0047 0.2510 6.4732

Coefficients:

Estimate Std. Error t value Pr(>|t|)

(Intercept) 0.0007587 0.0300654	0.025 0.97987

Line2	0.0096555 0.0425190	0.227 0.82036

Line3	-0.0566756 0.0425190 -1.333 0.18258

Line4	-0.0709358 0.0425190 -1.668 0.09528 .

Line5	0.0391742 0.0425190	0.921 0.35690

Line7	0.0011368 0.0425190	0.027 0.97867

Line8	0.0212880 0.0425190	0.501 0.61661

Line9	-0.0612518 0.0425190 -1.441 0.14974

Line11	-0.0078874 0.0425190 -0.186 0.85284

Line15	0.0088876 0.0425190	0.209 0.83443

Line28	0.0257850 0.0425190	0.606 0.54424

Line29	0.1008022 0.0425190	2.371 0.01777 *

Line50	0.0538299 0.0425190	1.266 0.20553

Line88	-0.0292602 0.0425190 -0.688 0.49136

Line108	-0.0778097 0.0425190 -1.830 0.06728 .

Line112	0.1257249 0.0425190	2.957 0.00311 **

Line115	0.0709096 0.0425190	1.668 0.09540 .

Line117	0.0520713 0.0425190	1.225 0.22073

Line119	-0.0103935 0.0425190 -0.244 0.80689

Line123	0.0261890 0.0425190	0.616 0.53795

Line152	-0.0404483 0.0425190 -0.951 0.34147

---

Signif. codes: 0 '***' 0.001 '**' 0.01 '*' 0.05 '.' 0.1 ' ' 1

Residual standard error: 0.6594 on 10080 degrees of freedom

Multiple R-squared: 0.006463,	Adjusted R-squared: 0.004492

F-statistic: 3.279 on 20 and 10080 DF, p-value: 1.001e-06

Call:

lm(formula = y ~ Line, data = chr14DataMA)

Residuals:

Min	1Q Median	3Q	Max

-7.8645 -0.2705 -0.0221 0.2515 6.3408

Coefficients:

Estimate Std. Error t value Pr(>|t|)

(Intercept) 0.028990	0.032410	0.894	0.371

Line2	0.022777	0.045835	0.497	0.619

Line3	-0.011204	0.045835 -0.244	0.807

Line4	-0.006384	0.045835 -0.139	0.889

Line5	0.042464	0.045835	0.926	0.354

Line7	0.050682	0.045835	1.106	0.269

Line8	0.016167	0.045835	0.353	0.724

Line9	0.504157	0.045835 10.999	<2e-16 ***

Line11	-0.004188	0.045835 -0.091	0.927

Line15	-0.002585	0.045835 -0.056	0.955

Line28	0.036347	0.045835	0.793	0.428

Line29	0.057268	0.045835	1.249	0.212

Line50	-0.002264	0.045835 -0.049	0.961

Line88	-0.005671	0.045835 -0.124	0.902

Line108	-0.007048	0.045835 -0.154	0.878

Line112	0.012519	0.045835	0.273	0.785

Line115	0.025095	0.045835	0.548	0.584

Line117	-0.039718	0.045835 -0.867	0.386

Line119	0.012036	0.045835	0.263	0.793

Line123	-0.023506	0.045835 -0.513	0.608

Line152	-0.043892	0.045835 -0.958	0.338

---

Signif. codes: 0 '***' 0.001 '**' 0.01 '*' 0.05 '.' 0.1 ' ' 1

Residual standard error: 0.661 on 8715 degrees of freedom

Multiple R-squared: 0.02659,	Adjusted R-squared: 0.02436

F-statistic: 11.91 on 20 and 8715 DF, p-value: < 2.2e-16

Call:

lm(formula = y ~ Line, data = chr15DataMA)

Residuals:

Min	1Q Median	3Q	Max

-8.8229 -0.2865 -0.0098 0.2702 6.7844

Coefficients:

Estimate Std. Error t value Pr(>|t|)

(Intercept) 0.034629	0.029034	1.193	0.2330

Line2	-0.032954	0.041060 -0.803	0.4222

Line3	-0.058301	0.041060 -1.420	0.1557

Line4	-0.036715	0.041060 -0.894	0.3712

Line5	0.054686	0.041060	1.332	0.1829

Line7	0.018381	0.041060	0.448	0.6544

Line8	0.003219	0.041060	0.078	0.9375

Line9	-0.091813	0.041060 -2.236	0.0254 *

Line11	-0.017415	0.041060 -0.424	0.6715

Line15	-0.027566	0.041060 -0.671	0.5020

Line28	0.004058	0.041060	0.099	0.9213

Line29	0.075875	0.041060	1.848	0.0646 .

Line50	-0.033880	0.041060 -0.825	0.4093

Line88	-0.030914	0.041060 -0.753	0.4515

Line108	-0.004641	0.041060 -0.113	0.9100

Line112	0.095591	0.041060	2.328	0.0199 *

Line115	0.011767	0.041060	0.287	0.7744

Line117	-0.025635	0.041060 -0.624	0.5324

Line119	-0.029526	0.041060 -0.719	0.4721

Line123	-0.063314	0.041060 -1.542	0.1231

Line152	-0.062302	0.041060 -1.517	0.1292

---

Signif. codes: 0 '***' 0.001 '**' 0.01 '*' 0.05 '.' 0.1 ' ' 1

Residual standard error: 0.692 on 11907 degrees of freedom

Multiple R-squared: 0.004192,	Adjusted R-squared: 0.002519

F-statistic: 2.506 on 20 and 11907 DF, p-value: 0.0002177

Call:

lm(formula = y ~ Line, data = chr16DataMA)

Residuals:

Min	1Q Median	3Q	Max

-5.8128 -0.2733 -0.0122 0.2534 8.1818

Coefficients:

Estimate Std. Error t value Pr(>|t|)

(Intercept) 0.059440	0.027832	2.136 0.032732 *

Line2	-0.044433	0.039361 -1.129 0.258985

Line3	-0.062596	0.039361 -1.590 0.111793

Line4	-0.088237	0.039361 -2.242 0.024999 *

Line5	0.018447	0.039361	0.469 0.639320

Line7	-0.025207	0.039361 -0.640 0.521926

Line8	-0.034723	0.039361 -0.882 0.377701

Line9	-0.123296	0.039361 -3.132 0.001739 **

Line11	-0.031360	0.039361 -0.797 0.425623

Line15	-0.042250	0.039361 -1.073 0.283112

Line28	-0.007897	0.039361 -0.201 0.840982

Line29	0.138458	0.039361	3.518 0.000437 ***

Line50	0.059020	0.039361	1.499 0.133782

Line88	-0.081872	0.039361 -2.080 0.037547 *

Line108	-0.116158	0.039361 -2.951 0.003174 **

Line112	0.661864	0.039361 16.815 < 2e-16 ***

Line115	0.103540	0.039361	2.631 0.008538 **

Line117	0.075854	0.039361	1.927 0.053989 .

Line119	-0.052582	0.039361 -1.336 0.181610

Line123	0.046593	0.039361	1.184 0.236541

Line152	-0.065598	0.039361 -1.667 0.095631 .

---

Signif. codes: 0 '***' 0.001 '**' 0.01 '*' 0.05 '.' 0.1 ' ' 1

Residual standard error: 0.6098 on 10059 degrees of freedom

Multiple R-squared: 0.06429,	Adjusted R-squared: 0.06243

F-statistic: 34.56 on 20 and 10059 DF, p-value: < 2.2e-16
